# Distinct sub-cellular autophagy impairments occur independently of protein aggregation in induced neurons from patients with Huntington’s disease

**DOI:** 10.1101/2021.03.01.433433

**Authors:** Karolina Pircs, Janelle Drouin-Ouellet, Jeovanis Gil, Melinda Rezeli, Daniela A. Grassi, Raquel Garza, Yogita Sharma, Isabelle St-Amour, Marie E. Jönsson, Pia A. Johansson, Kate Harris, Romina Vuono, Thomas Stoker, Bob A. Hersbach, Kritika Sharma, Jessica Lagerwall, Stina Lagerström, Petter Storm, Vivien Horváth, Sébastien S. Hébert, György Marko-Varga, Malin Parmar, Roger A. Barker, Johan Jakobsson

## Abstract

Huntington’s disease (HD) is a neurodegenerative disorder caused by CAG expansions in the huntingtin (HTT) gene. Modelling HD has remained challenging, as rodent and cellular models poorly recapitulate the disease. To address this, we generated induced neurons (iNs) through direct reprogramming of human skin fibroblasts, which retain age-dependent epigenetic characteristics. HD-iNs displayed profound deficits in autophagy, characterised by reduced transport of late autophagic structures from the neurites to the soma. The neurite-specific alterations in autophagy resulted in shorter, thinner and fewer neurites presented by HD-iNs. CRISPRi-mediated silencing of *HTT* did not rescue this phenotype but rather resulted in additional autophagy alterations in ctrl-iNs, highlighting the importance of wild type *HTT* in neuronal autophagy. In summary, our work identifies a distinct subcellular autophagy impairment in aged patient derived HD-neurons and provides a new rational for future development of autophagy activation therapies.

## Introduction

Huntington’s disease (HD) is an autosomal dominant neurodegenerative disorder caused by an expanded polyglutamine tract within the first exon of *Huntingtin* (*HTT*) (1). Clinically, HD is characterized by involuntary movements together with cognitive impairment, psychiatric disturbances as well as metabolic and sleep problems, a result of extensive cell impairment and death within the central nervous system (CNS). Genetics and age in combination are key components of HD pathology as the length of the CAG repeat expansion in *HTT* correlates with age of disease onset, and manifest disease is more prevalent with increased age, independent of CAG repeat length (1–3). Most HD patients have CAG repeats in the range of 40 - 45 CAGs and are diagnosed around the age of 50 (4). HTT is ubiquitously expressed, yet the presence of a mutated *Huntingtin* allele (m*HTT*) results, at least early on, in dysfunction and death of neurons specifically in the striatum and cortex (5). Mutant HTT has a propensity to aggregate and form insoluble protein inclusions, but it is still debated as to how protein aggregation influences, if at all, neuronal dysfunction and ultimately cell death. In general, the molecular and cellular basis for the pathology and the age-related disease process remains poorly understood and treatment of HD remains a major challenge.

Several studies have documented altered autophagy in neurodegenerative disorders including HD, a phenomenon thought to contribute to the failure of clearance of aggregating proteins (6–12). Autophagy is a lysosomal protein degradation pathway that is present at a basal level in all cells, including neurons and is essential for their survival (13, 14). Boosting autophagy through pharmacological or genetic manipulation successfully reverses disease-associated phenotypes in various mouse models of neurodegenerative disorders, including models of HD, and is associated with a reduction of the protein aggregate burden (6, 7, 11, 15, 16). These pre-clinical findings have led to the initiation of clinical trials to activate autophagy in HD and other neurodegenerative disorders (17–20). While these initial studies have shown that this approach is feasible and well tolerated, it is also evident that therapeutic approaches to activate autophagy need to be optimized and tailored for different neurodegenerative disorders. In particular, a clear understanding of exactly how and why alterations in autophagy appear in HD (and other neurodegenerative disorders) and how this contributes to neuronal dysfunction and death is currently lacking.

In this study we have used direct reprogramming of human fibroblasts to generate patient-derived induced neurons (iNs) that retain age-associated epigenetic marks (21–23). When performing a combined transcriptomic, proteomic and automated microscopic analysis on iNs obtained from patients with HD (HD-iNs), we found a distinct impairment of autophagy characterized by a failure to transport late autophagic structures from neurites to the cell body. This subcellular autophagy impairment was directly linked to a reduction in the neurite complexity of HD-iNs. The autophagy impairment in HD-neurons appeared without the presence of mHTT-aggregates, demonstrating that this phenomenon lies upstream of overt protein aggregation. Finally, inhibition of *HTT*-expression (both wt and mutant) using CRISPRi rescued some of the autophagy-related impairments but also resulted in additional new autophagy alterations suggesting that the disease phenotype is driven by a combination of both loss-of-function and gain-of-function mechanisms. In summary, our results provide a novel understanding of the HD disease process by demonstrating a specific subcellular autophagy impairment localised to the neurites. Our findings have clear translational implications.

## Results

### Direct iN conversion of fibroblasts from HD patients

We collected fibroblasts through skin biopsies from 10 individuals diagnosed with HD and 10 age- and sex-matched healthy controls (ctrl) (Table 1). The HD patients were all adult onset and between 28 - 59 years of age with CAG repeats lengths in the range between 39 – 58 (Table 1). The CAG-repeat length was initially determined by genotyping patient biopsies and later confirmed in the established fibroblast cultures. HD and control fibroblasts had a similar morphology and could be expanded at similar rate.

**Table 1.** Summary of control and HD patient biopsies. Overview of the cohort used in the study specifying the age, sex and CAG repeats of 10 healthy control and 10 HD patient fibroblasts lines.

| Line | Age | Sex | CAG repeats |
| --- | --- | --- | --- |
| C1 | 27 | M | 17/n/a |
| C2 | 30 | M | 19/24 |
| C3 | 52 | F | 19/23 |
| C4 | 54 | F | 15/20 |
| C5 | 61 | F | 17/n/a |
| C6 | 61 | M | 17/23 |
| C7 | 66 | M | 24/n/a |
| C8 | 67 | F | n/a |
| C9 | 71 | M | n/a |
| C10 | 75 | F | 18/n/a |
| HD1 | 28 | M | 15/39 |
| HD2 | 31 | M | 20/45 |
| HD3 | 33 | F | 17/58 |
| HD4 | 38 | F | 17/52 |
| HD5 | 43 | M | 17/42 |
| HD6 | 43 | M | 19/44 |
| HD7 | 47 | M | n/a/40 |
| HD8 | 49 | F | 18/47 |
| HD9 | 53 | M | 19/42 |
| HD10 | 59 | M | 16/39 |

We reprogrammed the 20 fibroblast lines to iNs using our previously described protocol (21, 24). In brief, this methodology includes a single lentiviral construct that expresses the transcription factors Achaete-scute homolog 1 (*Ascl1)* and POU Class 3 Homeobox 2 (*Pou3f2v* or *Brn2)* with two short hairpin RNAs (shRNA) targeting RE1-silencing transcription factor (REST1) (21) (Figure 1a). Upon transduction, the fibroblasts rapidly developed a clear neuronal morphology with a reduction in the size of both the nuclei and cell body and the formation of long, elaborate neurites (Figure 1b, SFigure 1a-b). Over a time period of a few weeks, the reprogrammed fibroblasts transformed into mature iNs and started to express the neuronal markers MAP2 (neuron specific cytoskeletal protein enriched in dendrites) and TAU (a highly soluble microtubule-associated protein abundant in neurons) (Figure 1b, SFigure 1a-b). In addition to which they became electrically active as we have previously shown (21).

**Figure 1.**
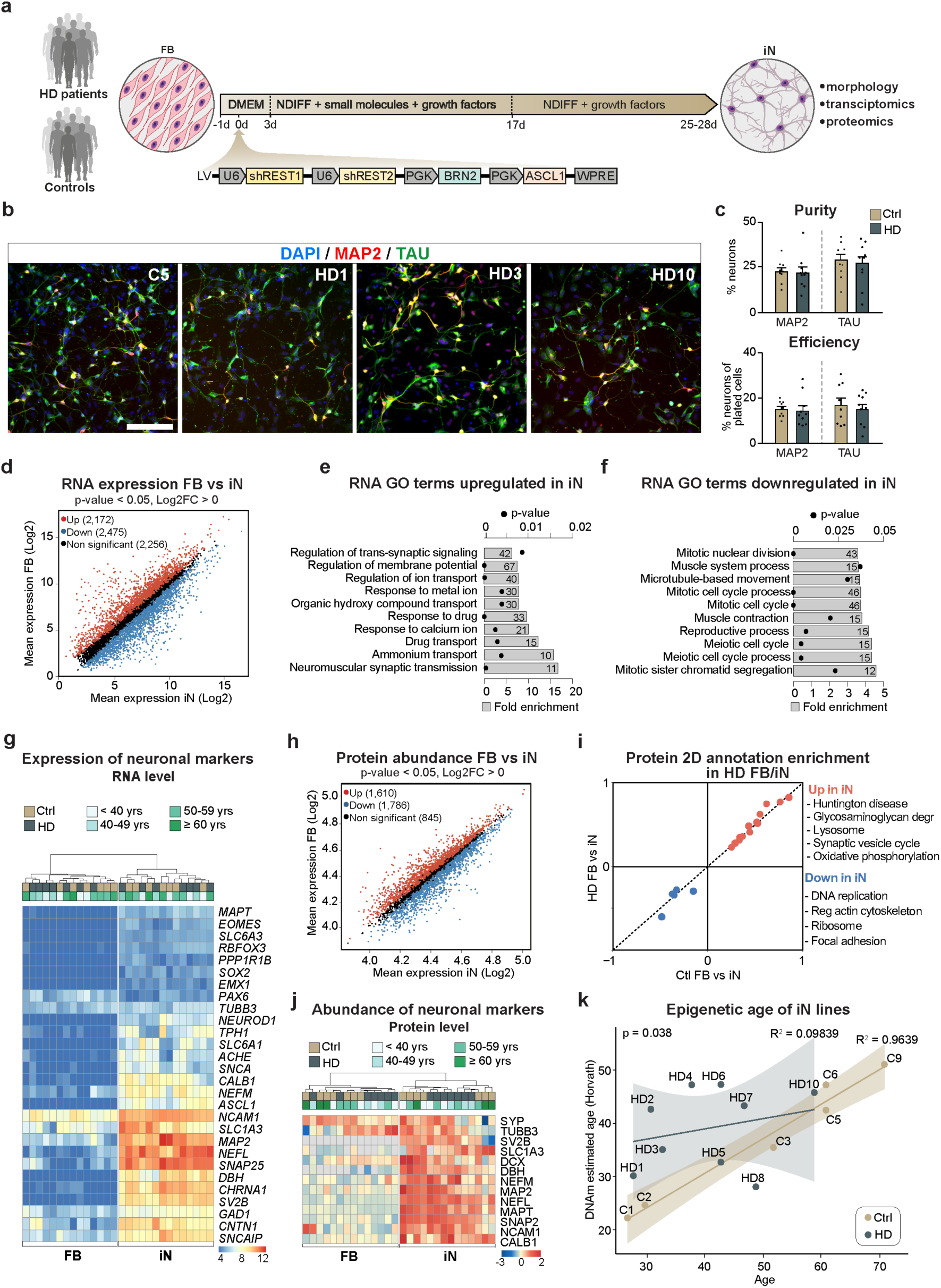
HD fibroblasts readily convert into iNs with similar purity and conversion efficiency. (a) Experimental overview of the iN conversion. (b) iNs derived from control and HD patient fibroblasts both express mature neuronal markers like TAU and MAP2. (c) Percentage of MAP2^+^ or TAU^+^ neurons from DAPI^+^ cells. Each dot represents the average value for one control or HD cell line. Percentage of MAP2^+^ or TAU^+^ neurons from plated cells after conversion (n = 9 lines for controls, 81 wells analyzed for MAP2 in total and 78 for TAU; n = 10 lines for HD, 85 wells analyzed in total for MAP2 and 77 for TAU). (d) Scatter plot displaying RNA-sequencing log2 mean gene expression in iNs (x-axis) and fibroblasts (y-axis). Significantly upregulated genes in iNs compared to fibroblasts are shown in red, significantly downregulated genes are shown in blue, and non-significant genes in black* (n = 7 control and 7 HD fibroblast and iN lines). (e-f) Gene ontology overrepresentation test of biological processes (Fisher’s Exact test using PANTHER GO-slim biological process) of genes up or downregulated in iNs compared to fibroblasts (Differential gene expression analysis performed with DESeq2; padj <0.05, log2FC >1), top ten most significant terms are shown. Grey bar plots represent fold enrichment. Circles represent Benjamini-Hochberg false discovery rates (n = 7 control and 7 HD fibroblast and iN lines; FDR<0.05). (g) Heat map of RNA expression of neural markers (n = 7 control and 7 HD fibroblast and iN lines; normalized by mean of ratios, padj <0.05). (h) Scatter plot displaying mean protein abundance in iNs (x-axis) and fibroblasts (y-axis). Proteins with statistically significant differences between groups were highlighted in red (upregulated in neurons) or blue (downregulated in neurons) *. Proteins that were not found significantly different are shown in black (n = 7 control and 7 HD fibroblasts and iN lines). (i) 2D annotation enrichment analysis of biological pathways between iNs and fibroblasts from HD patients and healthy donors. Significant pathways were selected following a threshold of 0.02 (Benjamini-Hochberg FDR). (j) Heat map of protein abundance of neural markers (n = 7 control and 7 HD fibroblast and iN lines; normalized counts, padj <0.05). (k) Scatter plot of chronological age in years (x-axis) versus DNAm predicted age (y-axis) with regression curves and 95%-confidence intervals plotted separately for control and HD-iNs (n = 6 for control and 9 HD-iN lines; Pearson correlation coefficient R^2^ = 0.9639 for control and 0.09839 for HD-iN lines). (*p<0.05; two-tailed unpaired T-tests were used) All data are shown as mean ± SEM. Scale bar is 50 μm. See also SFigure 1 and 2. FB: fibroblasts, GO: gene ontology, iN: human induced neurons, DMEM: Dulbecco’s modified eagle medium, Ndiff: Neural differentiation medium, sh: short hairpin, REST1/2: RE1/2-silencing transcription factor, PGK: Phosphoglycerate kinase promoter, BRN2: POU Class 3 Homeobox 2, ASCL1: Achaete-Scute Family BHLH Transcription Factor 1, WPRE: Woodchuck Hepatitis Virus Posttranscriptional Regulatory Element

We analyzed the reprogramming capacity of fibroblasts derived from HD-patients in detail using a high-content automated microscopy analysis of the reprogrammed iNs. By quantifying the number and proportion of MAP2^+^ and TAU^+^ cells (as defined by DAPI) we found that the HD fibroblasts converted into iNs four weeks post-transduction with similar purity (number of iNs / number of DAPI cells) and conversion efficiency (number of iNs / number of starting fibroblasts) (Figure 1c, SFigure 1c-d) as to that seen with control fibroblasts. Neuronal purity and conversion efficiency were not affected by passage number (SFigure 1e-f), and there was no difference in the rate of cell death between control and HD-iNs at four weeks, as determined through surveying the number of iNs and DAPI^+^ cells at this stage (SFigure 1g). Together, these data demonstrate that fibroblasts obtained from HD patients can be reprogrammed to iNs with the same efficiency as that seen for healthy control individuals.

### Transcriptome, proteome and epigenome profiling of iNs

To investigate molecular changes during the reprogramming process as well as molecular alterations in HD-iNs, we performed transcriptome and proteome analysis using RNA sequencing and shotgun proteomics on ctrl and HD-iNs, as well as the unconverted fibroblasts.

To obtain a pure population of iNs for these analyses (in order to reduce background transcriptional noise), we established a procedure to FACS-purify iNs at four weeks post conversion using Neural cell adhesion molecule (NCAM+), a mature neuronal cell surface marker (24) (Figure 1a).

We first analyzed the RNA-seq transcriptome data across seven ctrl and seven HD iNs and fibroblasts and found that fibroblast and iNs samples (both ctrl and HD) were clearly distinguishable (Figure 1d, SFigure 2a, STable 1, 2). We found high-level RNA expression of numerous genes that are known to be expressed in neurons in the iNs but not in the fibroblasts confirming successful neuronal conversion. Gene ontology analysis confirmed that transcripts enriched in iNs represented cellular processes related to neuronal functions, such as synaptic signaling and regulation of membrane potential (Figure 1e). On the other hand, transcripts enriched in fibroblasts were related to cell proliferation (Figure 1f). We also investigated the presence of transcripts related to specific neuronal subtypes in the iNs and found genes related to several different neuronal subtypes (Figure 1g), as well as an absence of neural progenitor markers both in the fibroblasts and in the iNs (Figure 1g). This is in line with previous results indicating that these types of iNs represent a mixed population of maturing neurons (21–23, 25).

**Figure 2.**
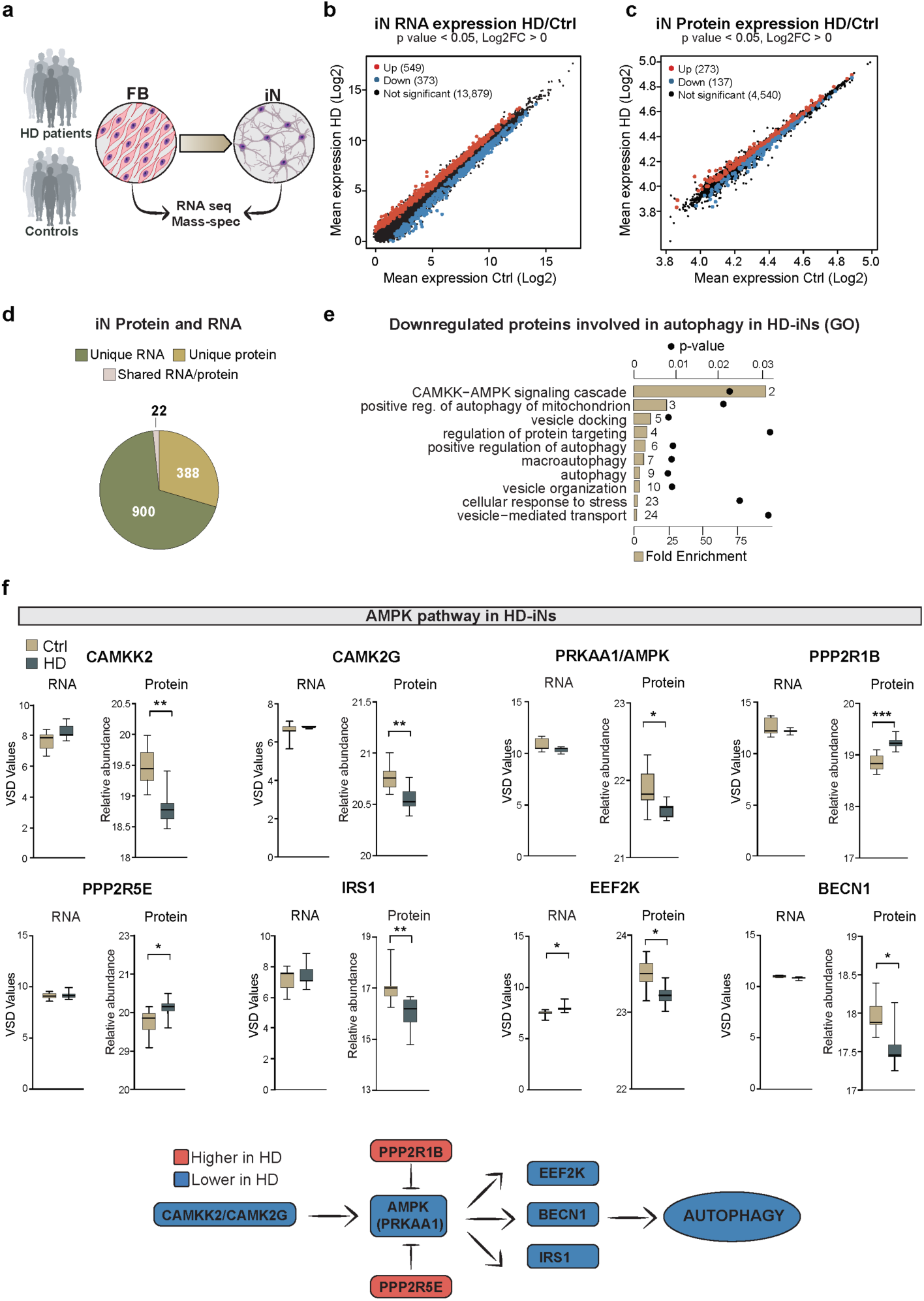
HD-iNs show a major post-transcriptional difference using quantitative proteomics. (a) Experimental overview of RNA-seq and Shotgun proteomic experiments. (b-c) Scatter plots displaying log2 mean gene expression or protein abundance in control and HD-iNs. Significantly upregulated RNAs and proteins in HD-iNs compared to controls are shown in red, downregulated RNAs/proteins in HD-iNs compared to controls are shown in blue, and non-significant genes in black* (n = 7 control and 7 HD-iN lines). (d) Number of significantly differentially expressed RNAs or proteins in control and HD-iNs. (e) Selected biological processes connected to autophagy by gene ontology functional enrichment analysis (STRING, biological process) of proteins downregulated in HD-iNs compared to ctrl-iNs. Grey bar plots represent fold enrichment. Circles represent P values (n = 7 control and 7 HD fibroblast and iN lines; p<0.05). (f) AMPK pathway proteins significantly dysregulated between control and HD-iNs where the RNA expression was not changed (n = 7 control and 7 HD-iN lines). (***p<0.001; **<0.01; *p<0.05; two-tailed unpaired T-tests were used in all) All data are shown as min/max box plots. See also SFigure 3.

Next, we analyzed the shotgun proteomics data from the unconverted seven ctrl and seven HD fibroblasts as well as the resulting iNs. The proteome analysis resulted in 7,001 proteins being quantified and identified at high confidence in the majority of samples in at least one group (Figure 1h, STable 3, 4). When we compared the abundance of individual proteins, we found that fibroblast samples and iNs (both ctrl and HD) displayed a high degree of proteomic difference when compared each other (SFigure 2b), similar to what was observed in the transcriptomic analysis. In particular, proteins linked to neuronal function, such as synaptic vesicles proteins, were highly abundant in iNs, while proteins related to proliferation pathways, such as cell cycle and DNA-replication were downregulated compared to fibroblasts (Figure 1i, SFigure 2c-e). Additionally, the metabolic reprogramming observed in iNs, involved the upregulation of pathways like glycolysis, the lysosome and phagosome, demonstrating that these cells, to a large extent, mimic the metabolic state normally found in neurons (Figure 1i-j, SFigure 2f-h).

Several previous studies have demonstrated that iNs retain age-dependent molecular features (22, 23, 26–29). To confirm this in our iNs, we investigated if we could detect age-dependent epigenetic signatures in these cells. We used the Illumina Epic Methylation array to profile global DNA methylation patterns in 6 ctrl and 9 HD-iNs. A penalized regression model using a set of 353 CpGs defining the biological age by the Horvath epigenetic clock allows the prediction of the age of the donor (30). We converted ctrl and HD donor cell lines into iNs and estimated the biological age of the resulting iNs. We found that in the ctrl-iNs, the DNAm predicted biological age strongly correlated with the donor’s actual real age (Pearson correlation coefficient R^2^ = 0.9639, Figure 1k). A previous study performed on postmortem brain tissue indicated an increase in epigenetic aging rates in patients with HD (31). In line with this, we also found significantly increased DNAm predicted biological age in the HD-iNs compared to the ctrl-iNs (p = 0.038, Figure 1k). Taken together, these data confirm that both ctrl and HD-iNs retain epigenetic signatures consistent with aged neuronal cells and that iNs derived from patients with HD have an increased biological age.

### HD-iNs display alterations in proteins linked to autophagy

To identify molecular mechanisms potentially linked to HD pathogenesis, we analyzed HD-iNs for differences in their transcriptome and proteome when compared to ctrl-iNs (Figure 2a). Starting with the transcriptome, we found 549 mRNA transcripts that were upregulated in HD-iNs and 373 downregulated out of 13,879 detected transcripts compared to ctrl-iNs, confirming previous findings that m*HTT* induces major transcriptional alterations (Figure 2b, STable 5, 6). However, gene ontology and network analysis failed to identify any molecular or biological processes that were significantly enriched in the differentially expressed genes (STable 7, 8). This suggests that while HD-iNs display transcriptome alterations, these alterations are not linked to distinct gene programs, making it difficult to link transcriptomic changes in HD-iNs to phenotypical alterations.

We next turned our attention to the the differences between the proteomes of ctrl-iNs and HD-iNs and found that 273 proteins were upregulated in HD-iNs while 137 proteins were downregulated out of 4,950 proteins detected (Figure 2c, STable 9, 10). Noteworthy, the majority of proteins altered in HD-iNs were not changed at the RNA-level. Out of the 410 proteins that were either up- or downregulated in HD-iNs only 22 of these genes were altered at the mRNA level (Figure 2d, STable 11). This suggests that only a very limited fraction of the differentially expressed transcripts that we detected resulted in significant changes at the protein level. Rather, the vast majority of changes at the protein level appear to be the result of post-transcriptional mechanisms.

Interestingly, when we performed gene ontology and network analysis of significantly dysregulated proteins in HD-iNs, we found that many of these proteins were functionally linked (STable 12, 13). Downregulated proteins were enriched for cellular pathways such as the CAMKK-AMPK-signaling cascade as well as autophagy related processes, while upregulated proteins were connected to ribosomal functions (Figure 2e, STable 12, 13). We also performed the same analysis in ctrl and HD fibroblasts and found that ribosomal proteins were also upregulated in the HD-fibroblasts, suggesting that translational alterations may be a ubiquitous downstream consequence of the presence of m*HTT* (STable 14) (32). On the other hand, the proteins related to CAMKK-AMPK-signaling and autophagy were only downregulated in HD-iNs and not in the corresponding HD-fibroblasts, indicating that these proteomic alterations are linked to neuron-specific cellular functions (SFigure 3, STable 15). We thus focused our further analyses on these neuron-specific proteome alterations.

**Figure 3.**
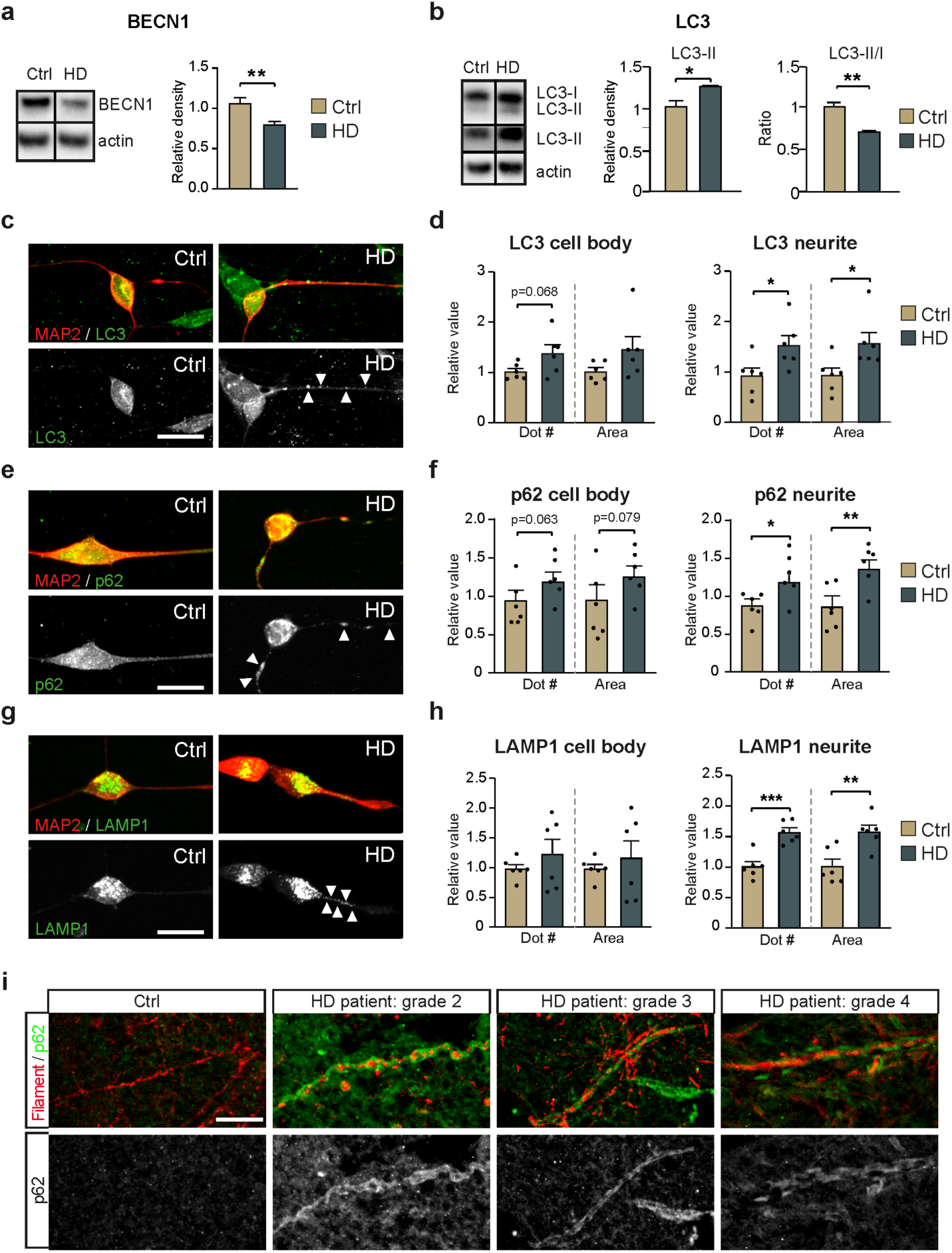
HD-iNs exhibit neurite specific autophagy alteration. (a) Reduced BECN1 expression in HD-iNs compared to ctrl-iNs using WB (n = 10 replicates for control and n = 9 replicates for HD-iNs). (b) LC3B-II levels are significantly increased in the HD-iNs, while the LC3B-II/I ratio decreased compared to the healthy ctrl-iNs (n = 6 replicates). (c-h) Representative images and statistical analysis shows a significant increase both in number and size of LC3B, p62 and LAMP1 dots in the MAP2^+^ neurites of HD-iNs compared to controls (n = 6 lines). (i) Representative images of human post-mortem striatal tissue from a healthy control and 3 different HD patients at different disease stages showing p62 accumulation specifically in the neurites as visualized by a neurofilament specific antibody. (***p<0.001; **<0.01; *p<0.05; two-tailed unpaired T-tests were used) All data are shown as mean ± SEM. WB values were normalized to ctrl-iNs expression levels and corrected to actin values. Scale bar is 25 μm. See also SFigure 4.

In HD-iNs, several kinases in the CAMKK-AMPK pathway were downregulated, including CAMKK2, CAMK2G, AMPK and IRS1 as well as the autophagy regulator BECN1 (Figure 2f). Moreover, suppressors of the AMPK pathway, PPP2R5E and PPP2R1B phosphatases were significantly upregulated in HD-iNs compared to healthy controls (Figure 2f). Taken together, this omics-based analysis demonstrate that HD-iNs display an altered proteome with links to alterations in autophagy. Noteworthy, these alterations were cell-type specific, only present in the iNs but not in the fibroblasts and mainly due to post-transcriptional mechanisms that could not be detected by transcriptome analysis.

### Subcellular alterations in autophagy in HD-iNs

One of the downregulated proteins detected in the proteomic analysis was BECN1, an autophagic regulator protein that plays a key role in autophagosome formation. Several studies support the importance of BECN1 in HD pathology, as overexpression can slow the progression of HD pathology in both cell and mouse models by inducing autophagy, while the expression of BECN1 in the brains of HD patients declines with age (6, 7, 33, 34). The downregulation of BECN1, as well as the other alterations in the CAMKK-AMPK-signaling pathway, suggests that autophagy activity may be impaired in HD-iNs.

To investigate this in more detail, we first verified that in HD-iNs (but not the parental fibroblasts) there was a significant reduction of BECN1 levels using western blot (WB) analysis (Figure 3a, SFigure 4a). We then assessed autophagy activity in HD-iNs compared to ctrl-iNs by measuring microtubule-associated protein 1A/1B-light chain 3B (LC3B). LC3B conjugates from LC3B-I to LC3B-II during autophagosome formation. We found a reduction in the LC3B conjugation, as determined by assessing the ratio of LC3B-II over LC3B-I using WB in HD-iNs (Figure 3b) which was coupled to an increase in total LC3B-II levels, suggesting more autophagosomes in HD-iNs (Figure 3b). We next measured p62 levels, which inversely reflects autophagolysosome degradation (35) and found a non-significant trend for increased levels (SFigure 4b). We also measured the levels of LAMP1, which is present on endosomes and lysosomes including autophagolysosomes and autolysosomes but did not detected significant difference in HD-iNs (SFigure 4b) (36). In summary, there were alterations in basal autophagy in HD-iNs, primarily reflected by alterations in BECN1 and LC3B, in line with an increase in autophagosomes and most likely a reduction in autophagic flux.

**Figure 4.**
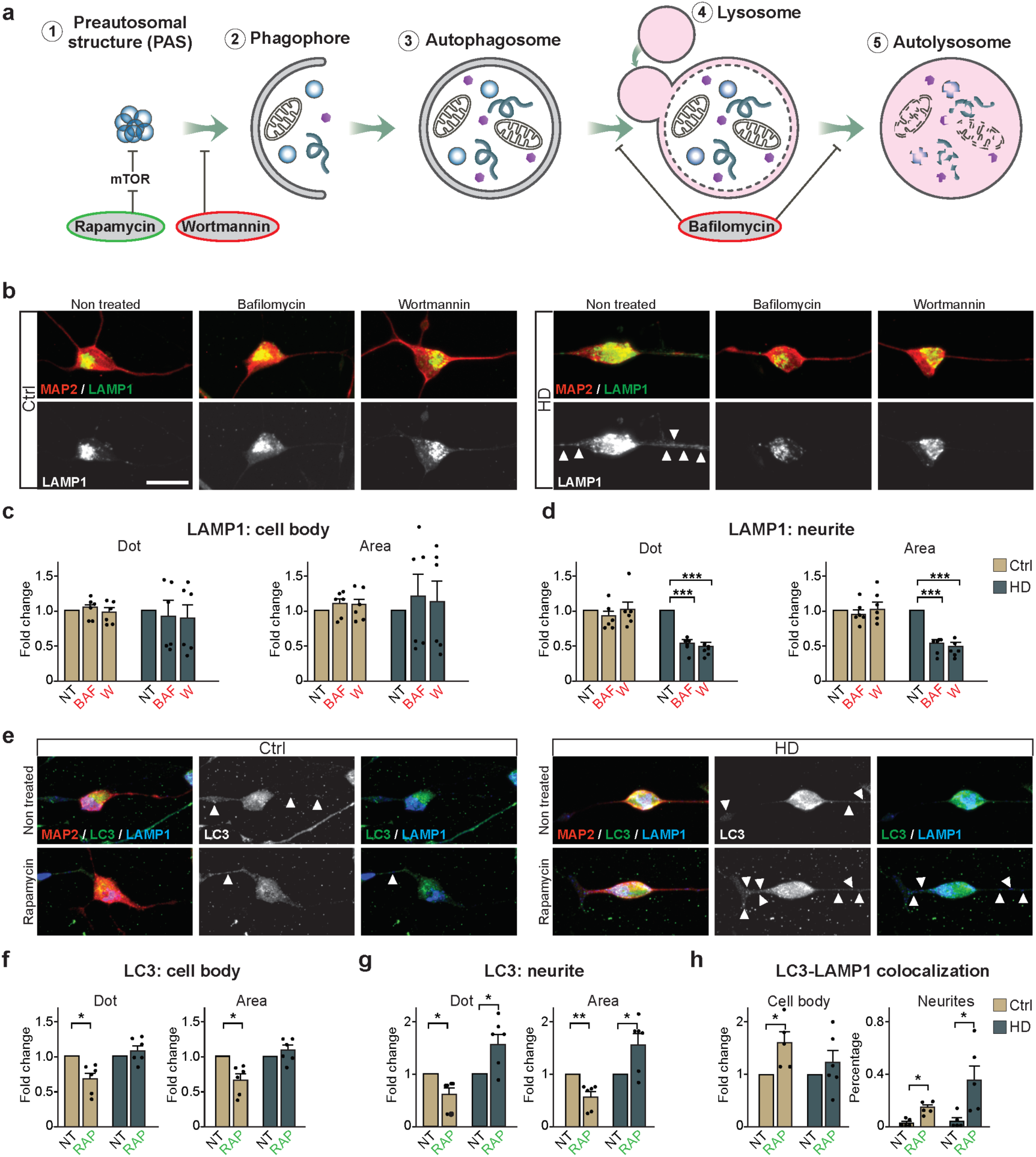
Autophagic flux is altered in the neurites in HD-iNs. (a) Schematic summary of the effect of different autophagy drugs. (b-d) Representative images and fold changes summarizing LAMP1^+^ dot number and area changes in the cell body and neurites of non-treated and Baf or W treated healthy control and HD-iNs (n = 6 lines). (e-g) Representative images of non-treated and rapamycin-treated healthy control and HD-iNs stained with the neuronal marker MAP2 together with LC3B and LAMP1. Arrowheads are indicating LC3, p62, LAMP1 positive dots in the neurites. Statistical analysis shows a significant decrease after RAP treatment both in the number and size of LC3B dots in the MAP2^+^ cell bodies and neurites in the control iNs (n = 6 lines). Statistical analysis shows an opposing effect of RAP treatment regarding the amount and area of LC3B puncta in the neurites between control and HD-iNs. While in the control iNs rapamycin significantly decreased the number and size of LC3B positive dots in the MAP2^+^ neurites, HD-iNs exhibited the opposite, LC3B dots significantly increased both in number and size (n = 6 lines). (h) Statistical analysis showing a significant increase in LC3B-LAMP1 colocalization in the cell bodies of ctrl-iNs, while there was no change in the HD-iNs (n = 6 lines). The percentage of LC3B-LAMP1 colocalization significantly increased both in the control and HD-iN neurites (n = 6 lines). (***p<0.001; **<0.01; *p<0.05; two-tailed paired T-tests were used in almost all cases except h (neurites panel) where one-way ANOVA was used) All data are shown as mean ± SEM. Fold changes are presented, except in figure h, neurite colocalization, where several datapoints were 0 therefore the colocalization is presented as percentage between LC3B and LAMP1. Scale bar is 25 μm. See also SFigure 5.

In neurons, autophagosomes are formed in the neurites and then transported to the cell body where the active lysosomes are present (37). Moreover, degradative lysosomes in the soma can also be transported to target autophagosomes in the distal axons anterogradely in mature neurons (38). To characterise autophagy alterations at a subcellular level, we performed immunocytochemistry (ICC) analysis of LC3B, p62 and LAMP1 as well as an unbiased quantification of autophagosomes including the subcellular localization using high-content automated microscopy. This analysis revealed an increased number and size of LC3B puncta in HD-iNs, that was particularly apparent in the neurites of these cells (Figure 3c, d). This demonstrates that autophagosomes accumulate specifically in neurites in HD-iNs. The increase in autophagosomes was coupled to an increased number and size of p62 puncta in neurites as well as an increase in the number and size of LAMP1-positive puncta at this same location (Figure 3e-h), indicating that autophagosomes and autophagolysosomes remain in the neurites and fail to transport their cargo to the soma for degradation in HD-iNs.

We used immunohistochemistry (IHC) to verify the neurite-specific impairment of basal autophagy in HD neurons using human post-mortem brain tissue (Table 2). We found no evidence of accumulation of p62 positive dots in the neurites of healthy controls identified by co-labelling with a Neurofilament specific antibody (Figure 3i). In contrast, we found clear p62 accumulation in the neurites in all HD patients analysed regardless of disease stage (Figure 3i). Taken as a whole we therefore conclude that there is a subcellular, neurite specific autophagy alteration in HD neurons.

**Table 2.** Human samples. Related to Figure 3.

| Figure 3i | Brainbank ID | Age of death | Pathological Grade | Number of CAG repeats |
| --- | --- | --- | --- | --- |
| Ctrl | PT89 | 66 | - | - |
| HD patient:<br>grade 2 | H721 | 61 | 2 | 46 |
| HD patient:<br>grade 3 | H715 | 57 | 3 | 47 |
| HD patient:<br>grade 4 | H693 | 43 | 4 | 51 |

### Impaired autophagic flux in HD-iNs

We next focused on analysing autophagic flux in HD-iNs by modulating the autophagy pathway using pharmacological agents (Figure 4a). Treatment with Bafilomycin A1 (Baf), a late-stage inhibitor of autophagy that blocks autophagosome-lysosome fusion, resulted in the expected increase in size of autophagosomes as visualized by LC3B-puncta, in the cell body of both ctrl and HD-iNs (SFigure 5a-c). However, the number of LC3B dots increased significantly in the ctrl-iNs, but not in the HD-iNs. Furthermore, there was an increase in p62-puncta count in both the cell body and neurites in HD-iNs but not in the ctrl-iNs (SFigure 5d-f). Thus, blocking autophagolysosomal formation in HD-iNs resulted in a further reduction in autophagy activity, suggesting that degradation of these structures occurs at a reduced rate in HD-iNs. In line with this observation, we found that the accumulation of autophagolysosomes in HD-iN neurites, as visualised by LAMP1-puncta, was completely abolished upon Baf-treatment (Figure 4b-d). Thus, when the formation of new autophagolysosomes is prevented in HD-iNs the cells are capable of dealing with the accumulation of these structures in the neurites. We further corroborated these results by treatment with Wortmannin (W), which blocks the initiation of autophagy by inhibiting phosphatidylinositol 3-kinase (PI3K) (Figure 4a) (36), which showed that there was a robust reduction of LAMP1-puncta in the neurites of HD-iNs with this treatment (Figure 4b-d).

**Figure 5.**
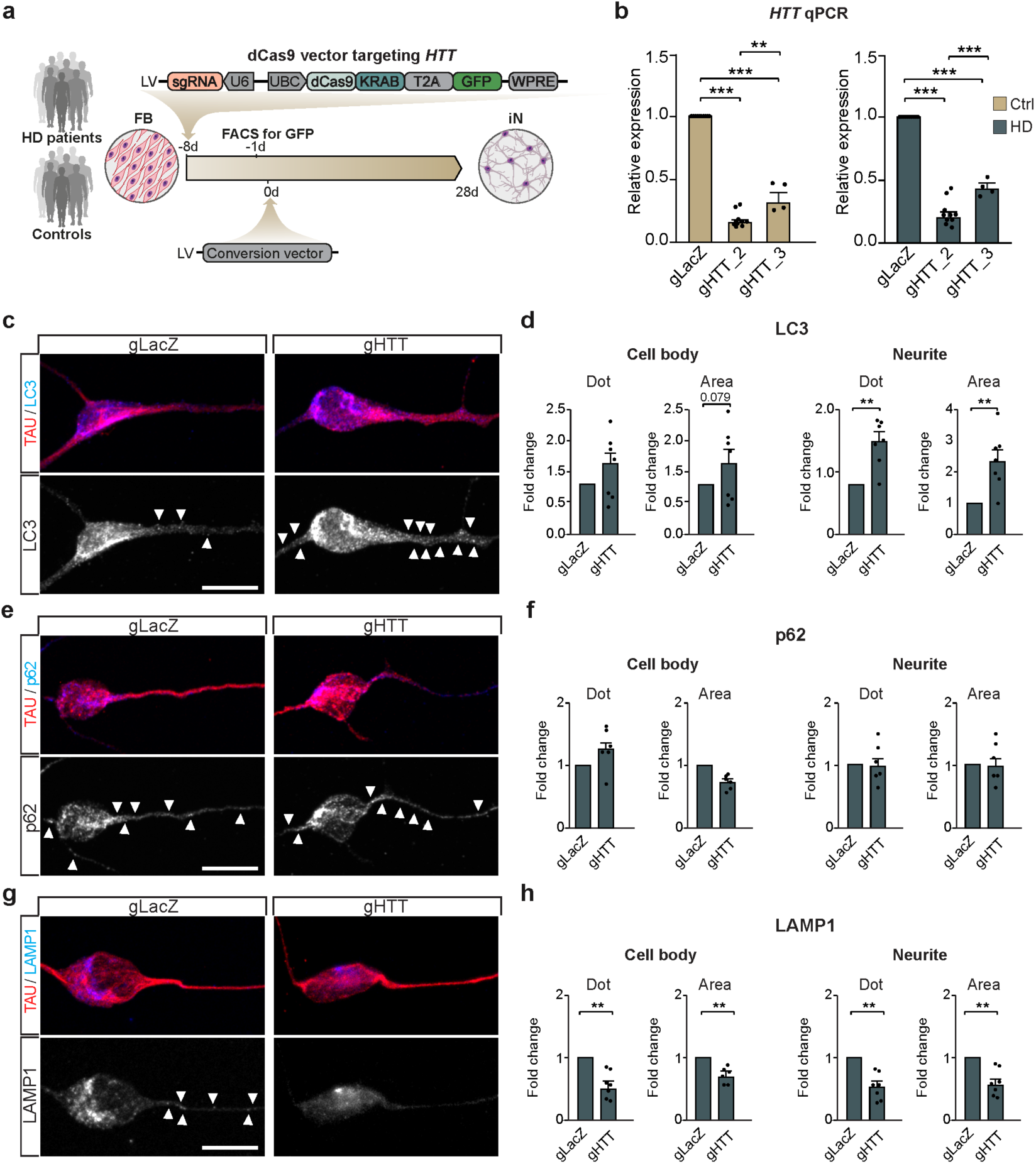
Silencing of HTT using CRISPRi further alters autophagy in HD-iNs. (a) Experimental overview. Fibroblasts from five HD patients and five healthy individuals were first transduced with lentiviral vectors targeting LacZ or HTT (sgRNA). After 7 days, GFP^+^ cells were FACS sorted and converted into iNs. (b) qRT-PCR revealed an efficient silencing of HTT using gRNA2 and gRNA3 both in control and HD-iNs (n = 10 replicates from 5 ctrl and 5 HD-iN lines for LacZ and gRNA2 and n = 4 replicates from 2 ctrl and HD-iN lines for gRNA3). (c-h) Representative images and statistical analysis of LC3, p62 and LAMP1 dot number and area in TAU^+^ cells in HD-iNs stably expressing LacZ and HTT gRNAs using CRISPRi (n = 7 replicates from 5 ctrl and 5 HD-iN lines pooling gRNA2 and gRNA3 data). Arrowheads are indicating LC3, p62, LAMP1 positive dots in the neurites. (**<0.01; *p<0.05; two-tailed paired T-tests were used) All data are shown as mean ± SEM. Fold changes are presented in all graphs. Scale bar is 25 μm. FB: fibroblasts, iN: human induced neurons, DMEM: Dulbecco’s modified eagle medium, Ndiff: Neural differentiation medium, sh: short hairpin, REST1/2: RE1/2-silencing transcription factor, PGK: Phosphoglycerate kinase promoter, BRN2: POU Class 3 Homeobox 2, ASCL1: Achaete-Scute Family BHLH Transcription Factor 1, WPRE: Woodchuck Hepatitis Virus Posttranscriptional Regulatory Element, UbC: mammalian ubiquitinC promoter, KRAB: Krüppel associated box transcriptional repression domain, T2A: thosea asigna virus 2A self-cleaving peptides. See also SFigure 6-8.

To further understand the autophagy impairment in HD-iNs we next used rapamycin (RAP) that activates autophagy at an early stage by inhibiting mTOR signalling (Figure 4a). In ctrl-iNs, RAP treatment resulted in a clear reduction in LC3B-puncta in both the cell body and neurites, in line with the increased autophagic flux mediated by the treatment (Figure 4e-g). However, in HD-iNs RAP treatment resulted in an increase in both the size and number of LC3B-puncta specifically in neurites (Figure 4e-g). Thus, the impairment in autophagolysosome transfer and degradation that is present in HD-iNs prevents an increased autophagic flux in RAP treated cells. Moreover, LAMP1 dot number and area was significantly reduced after RAP treatment in the ctrl-iNs cell body where the active lysosomes are present and where the late autophagic structures are transported for degradation, while this was not seen in the HD-iNs. This further corroborates an autophagosomal transport failure in the HD-iNs. These results were verified by performing co-localization analysis for LC3B and LAMP1 (Figure 4e, h, SFigure 5g, h). While RAP clearly increased the autophagy flux in the cell body of ctrl-iNs by an increased formation of LC3B-LAMP1 double positive late autophagy structures, we did not detect these structures in HD-iNs (Figure 4e, h). On the contrary, HD-iNs exhibited a significant increase only in neurite LC3B-LAMP1 co-localization after RAP treatment (Figure 4e, h). This further verifies that while autophagolysosomes are formed in HD-iNs they fail to get degraded and transported to the cell body. Early activation of autophagy using RAP thus increases the amount of trapped autophagolysosomes in the neurites of HD-iNs.

Taken as a whole, these results demonstrate that HD-iNs show impairment in degrading autophagolysosomes. It appears that the cellular machinery is working at reduced rate and cannot degrade autophagy cargo, resulting in an accumulation in late stage autophagic structures. The reason for this impairment is likely to relate to the late autophagic structures getting stuck in the neurites and failing to be transported to the cell body where they should be degraded. This impairment in the last step of autophagy results in an overall reduction in autophagy activity. These observations are important from a therapeutic point of view as treatment paradigms to restore autophagy alterations in HD should aim to enhance autophagolysosome transfer and degradation rather than activating autophagy at an early stage, which could actually worsen the pathology.

### Cellular mechanisms underlying the autophagy impairments found in HD-iNs

We next investigated the molecular mechanisms underlying the autophagy impairment in HD-iNs. It has been suggested that protein aggregates are the key driver of the autophagy phenotype observed in neurodegenerative disorders. For example, long-term exposure to protein aggregates could eventually exhaust the autophagy machinery (39, 40). On the contrary, HTT has also been suggested to be directly linked to the cellular signalling pathway that controls autophagic activity (41, 42). HTT has been reported to directly bind to BECN1 via its polyQ-tract and modulation of this binding, either by loss of wtHTT levels or by the presence of an expanded polyQ-tract in mHTT, results in a reduction in BECN1 levels and an overall reduction in autophagic activity (7, 34). wtHTT has also been reported to directly interact with p62 to facilitate cargo engulfment in autophagy, indicating that the loss-of-function of one wild-type allele of *HTT* in HD may impair autophagy (42).

To investigate this further, we first looked for the presence of mHTT-aggregates in HD-iNs. Through the use of WB analysis with several different lysis conditions, we did not detect the presence of any m*HTT*-containing aggregates (SFigure 6a, b) even though the expression of *HTT*-mRNA in fibroblasts and iNs was similar in both groups (SFigure 6c). Together, these experiments demonstrate that the autophagy impairments in HD-iNs are present without evidence for overt HTT aggregation.

**Figure 6.**
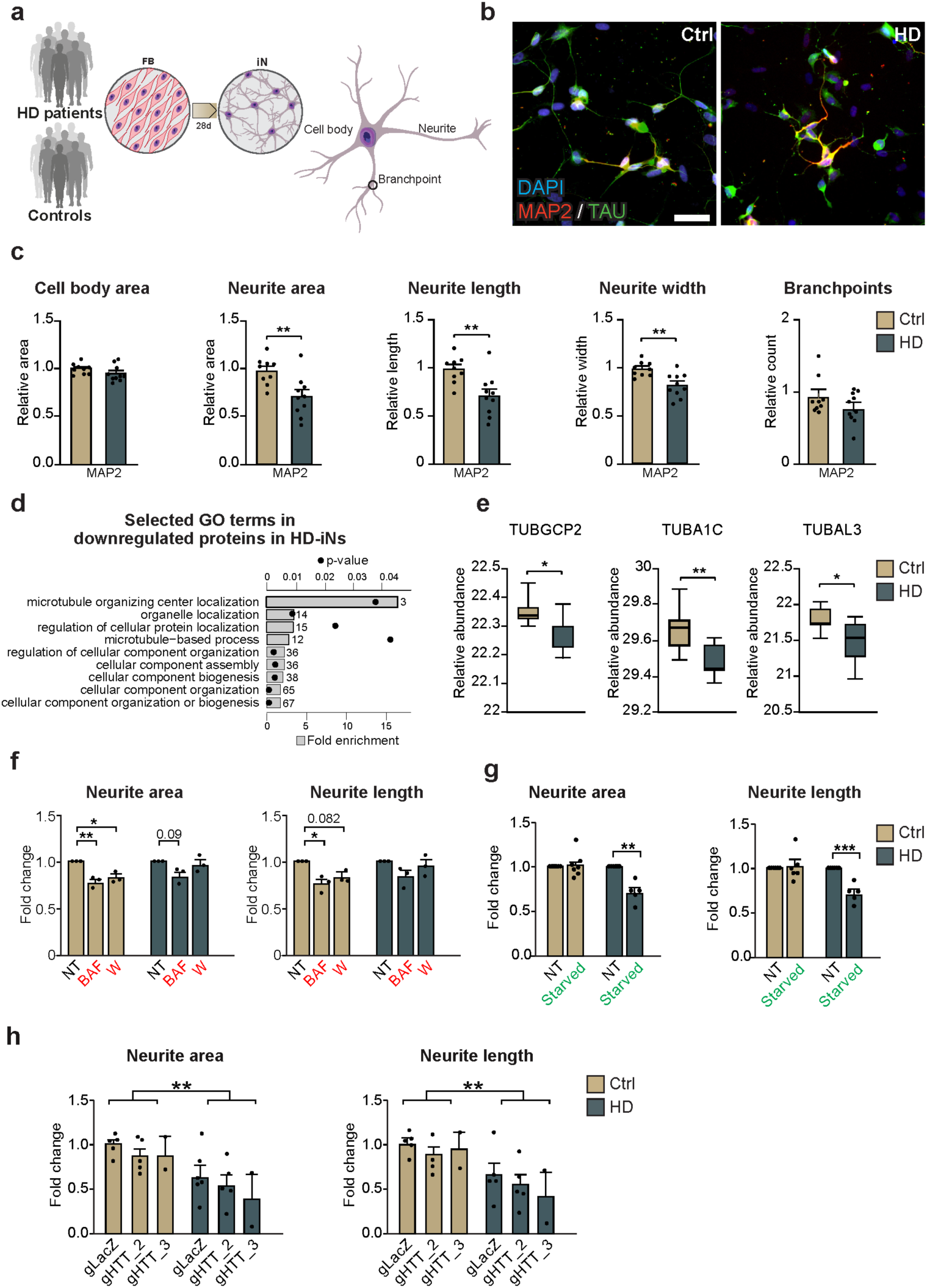
HD-iNs show a less elaborate neuronal morphology. (a) Experimental workflow summarizing iN conversion. After neural conversion, morphology of the cells is analyzed using high-content automated microscopy analysis. (b) Representative images after 28 days of conversion showing control and HD-iNs expressing mature neuronal markers like MAP2 and TAU. (c) The average relative cell body area and number of branchpoints per cells as defined by MAP2 staining using high-content automated microscopy analysis shows no difference between control and HD-iNs. Relative neurite area, length and width per cell was significantly reduced in the HD-iNs compared to the healthy controls (n = 9 lines for controls, 96 wells analyzed in total; n = 10 lines for HD, 119 wells analyzed in total). (d) Biological processes connected to microtubules and cytoskeletal organization selected from the gene ontology functional enrichment analysis (STRING, biological process) of proteins downregulated in HD-iNs compared to ctrl-iNs. Grey bar plots represent fold enrichment. Circles represent P values (n = 7 control and 7 HD fibroblast and iN lines; p<0.05). (e) Tubulin proteins significantly dysregulated between control and HD-iNs (n = 7 control and 7 HD-iN lines). (f) Neurite area and length per cell is reduced after autophagy impairment in control iNs, while it is not further reduced in HD-iNs (n = 3 control and 3 HD-iN lines, 9 – 9 wells analyzed in each condition). (g) Neurite area and length per cell is reduced after starvation in HD-iNs, while it is not changed in control iNs (n = 6 for ctrl-iN lines and n = 5 for HD-iN lines, 12 wells analyzed in total for control-iNs and 10 for HD-iNs). (h) Relative neurite area and length per cells were not changed in the HTT (wt and mutant) silenced HD-iNs compared to the LacZ transduced. HTT silencing did not affect neurite area and length in the control iNs (n = 5 ctrl and 5 HD-iN lines for LacZ and gRNA2, n = 2 ctrl and 2 HD-iN lines for gRNA3). (***p<0.001; **p<0.01; *p<0.05; two-tailed unpaired T-tests were used in c, e and g. Ordinary one-way ANOVA was used in f. Two-way ANOVA was used in h.) All data are shown as mean ± SEM in c and f-h. All data are shown as min/max box plots in e. Scale bar is 50 μm. See also SFigure 9.

We next investigated the direct role of *HTT* in the regulation of autophagy in iNs. Previous studies showed that silencing *HTT* blocks retrograde transport of late autophagosomes, while depletion of the m*HTT* results in accumulation of late autophagic structures with undegraded cargo (9, 10). Moreover, HTT is also involved in lysosomal transport (43, 44). Since both wild-type and mutant *HTT* have been implicated in the regulation of autophagy we decided to investigate the consequence of transcriptional silencing of wt*HTT*/m*HTT* on the autophagy pathway in both ctrl-iNs and HD-iNs (9). To this end we established a lentiviral based CRISPR inhibition (CRISPRi) approach to silence HTT-expression (Figure 5a). The CRISPRi-vector expressed a dead Cas9-KRAB fusion protein that was linked to a GFP reporter as well as a guide RNA (gRNA) targeted to the area around the *HTT* transcription start site (Figure 5a). This vector design allows for the binding of dCas9-KRAB to the *HTT* loci, thereby resulting in the establishment of local heterochromatin and subsequent transcriptional silencing. We optimized the vector construct by testing different gRNAs and MOIs in HEK293T cells and human induced pluripotent stem cells (iPSCs) and ultimately found two different gRNAs, targeted to a region just downstream of the *HTT* transcription start site (TSS), that very efficiently silenced both alleles of *HTT* (SFigure 7a).

We transduced ctrl and HD-fibroblasts with the CRISPRi-*HTT* vector and FACS purified GFP expressing cells (Figure 5a). This resulted in efficient silencing of both alleles of *HTT* in the patient-derived fibroblasts as quantified with qRT-PCR (SFigure 7b). We then proceeded to generate iNs from the CRISPRi-*HTT* silenced fibroblasts (Figure 5a). After four weeks of conversion, we confirmed that *HTT* remained silenced in the iNs after conversion and that CRISPRi-*HTT* treatment did not impact on reprogramming efficacy (Figure 5b, SFigure 7c-f). The resulting *HTT*-silenced HD-iNs and ctrl-iNs were then analyzed using ICC for LC3B, p62 and LAMP1 spots in the cell body and in the neurites.

We focused first on silencing of *HTT* in the ctrl-iNs. Previous studies have demonstrated that HTT has an essential function in autophagy, as it contains an autophagy-inducing domain and it also facilitates axonal trafficking of autophagosomes (9, 10). Moreover, HTT functions as a scaffold in autophagy where it physically interacts with p62 and depletion of HTT reduces the association of p62 with LC3B and other substrates of autophagy (42). When silencing *HTT* in ctrl-iNs we found that while LC3B dot number count or area were not affected, the number of p62 positive puncta significantly increased in the neurites of ctrl-iNs, confirming its role in regulating autophagy or other mechanisms related to p62 degradation (SFigure 8a, b). Notably, the number and area of LAMP1 puncta significantly decreased but only in the neurites of *HTT* silenced ctrl-iNs (SFigure 8c). Thus, silencing of *HTT* in the ctrl-iNs resulted in the alteration of autophagic activity characterised by increased p62 accumulation and reduction in the endolysosomal marker LAMP1. These findings are in line with previous studies demonstrating that HTT facilitates cargo recognition by modulating the assembly of the cargo receptors and autophagy proteins. Moreover, these findings highlight that silencing *HTT* in the ctrl-iNs results in a different autophagy impairment to that found in HD-iNs.

Next, we focused on the effect of silencing *HTT* in HD-iNs on autophagy. As described above it is important to highlight that CRISPRi experiments resulted in a highly efficient silencing of both healthy and m*HTT* alleles in the HD-iNs (Figure 5b). Moreover, as described above, HD-iNs display a neurite specific late-stage autophagy alteration with increased LC3B, p62, LAMP1 dot number and area. When silencing *HTT* (both the wt*HTT* and m*HTT* allele) in HD-iNs we found a further accumulation of LC3B both in terms of the number and their size in the neurites, while p62 expression was not significantly affected (Figure 5c-f). LAMP1 was significantly reduced in the HD-iNs after silencing *HTT* both in the neurites and in the cell body (Figure 5g, h). These results suggest that some of the autophagy impairments are restored by silencing m*HTT*, most notably there is a significant reduction of LAMP1 in the neurites. However, with this silencing comes another type of autophagic impairment likely due to a loss-of-function of the wild-type allele (Figure 5c-h, SFigure 8). Thus, CRISPRi silencing of wt*HTT*/m*HTT* does not substantially rescue the autophagy impairment in HD-iNs, most likely due to the important role of wt*HTT* in the control of autophagy.

### The autophagy impairment in HD-iNs results in reduction in neurite complexity

We finally explored the cellular consequences of the impaired autophagy in HD-iNs. It is well established that HD neurons tend to display alterations in neurite arborization and complexity, and these impairments are thought to contribute to the early disease process (26, 45–47). Importantly, autophagy has been directly linked to neurite formation, since inhibition of this degradation pathway reduces neurite growth and branching complexity (37, 48). To investigate whether HD-iNs have an altered neurite morphology and if this is linked to the autophagy impairments found in the cells, we performed a detailed analysis of neural morphology of the reprogrammed cells using high-content automated microscopy (Figure 6a, SFigure 9a, b). After four weeks of conversion, we found a significant decrease in neurite complexity in HD-iNs as measured by total neurite area, the number of neurites per cell, neurite length and neurite width (Figure 6b, c, SFigure 9c-f). This phenotype was not a consequence of a slower maturation of HD-iNs, since we observed a similar reduction in neurite number, length and complexity after seven weeks of conversion (SFigure 9g-m). Also, the cell body size was similar in HD-iNs and ctrl-iNs (Figure 6c, SFigure 9m). At a molecular level we found that many proteins that were downregulated in HD-iNs were connected to the microtubule system, which plays a fundamental role in the maintenance of axonal homeostasis by preserving axonal morphology and providing tracks for protein and organelle transport. A significant reduction was seen in proteins belonging to the tubulin protein superfamily, such as TUBGCP2, TUBA1C, TUBAL3, which are all involved in neuronal microtubule migration, axonal assembly and neurodegeneration (Figure 6d, e). Notably, these alterations in the microtubule system were dysregulated at a posttranscriptional level as the tubulin superfamily protein members were not different at the RNA level (SFigure 9n).

Autophagosomes form at the axon terminal and fuse with lysosomes during a dynein-mediated transport to the soma. Moreover, lysosome transport is also mediated via microtubules in the neurites (49). To investigate a direct link between the reduced neurite morphology in HD-iNs and autophagy, we analyzed neuronal morphology after inhibition or activation of autophagy using Baf or W and starvation, respectively. We found a significant reduction in the ctrl-iNs neurite area and length when inhibiting autophagy using Baf or W (Figure 7f). In contrast, HD-iNs did not exhibit any further reduction in neurite area or length after autophagy suppression (Figure 6f). These data suggest that while ctrl-iNs neurite morphology is affected by autophagy impairment using different pharmacological agents, HD-iNs do not show any further morphological changes. likely due to an already existing autophagolysosomal transport failure.

Next, we used amino acid free starvation to activate autophagy in the iNs. In response to starvation, cells recover nutrients through autophagy by increased AMPK activation and increased mTOR inhibition. This short-term autophagy activation through starvation did not have any major effect on the neuronal morphology of the ctrl-iNs since neurite area and length were not affected (Figure 6g). On the other hand, the neuronal morphology of HD-iNs was significantly affected, neurite area and length significantly decreased after starvation (Figure 6g), suggesting that HD-iNs could not cope even with this short-term starvation activation of autophagy. Lastly, we analyzed the effect of CRISPRi editing on the neurite morphology after silencing of *HTT* expression in ctrl-iNs and HD-iNs. CRISPRi silencing did not rescue the reduced neurite area nor neurite length in the HD-iNs (Figure 6h). HD-iNs were significantly shorter and smaller even after silencing both *HTT* alleles in the HD-iNs compared to the ctrl-iNs (Figure 6h). Together, these results suggest that the abnormal neuronal morphology present in the HD-iNs is directly linked to impairments in autophagy.

## Discussion

The pathogenic processes underlying HD have been difficult to elucidate, in part due to the fact that age-dependent human neurodegenerative disorders are challenging to study. Postmortem material is limited, both in availability and experimental possibilities and provides only a static snapshot of the consequence of disease. Animal models poorly replicate the disease process, as their lifespan is shorter in comparison to humans. This has led to the use of transgenic m*HTT*-alleles with very long CAG repeats (sometimes >100 CAGs), where the pathology is accelerated and thus possible to study in mice. However, this many repeats are rarely, if ever, seen in clinical practice. In case they are seen, they are associated with the rare juvenile form of the disease, in which the disease process may be drastically different from HD associated with more typical CAG repeat lengths (50). These challenges to model HD have contributed in part to the lack of effective treatments and it is therefore critical to establish model systems that recapitulate the human disease progression, including age-dependent processes.

Recent advances in cellular reprogramming have allowed for the establishment of induced pluripotent stem cells (iPSCs) that can be efficiently differentiated into neurons, making it possible to obtain human patient-derived HD-neurons (51–54) with the potential to generate isogenic control lines. However, a drawback with iPSCs is that during the reprogramming process epigenetic marks associated with ageing are erased, thereby transforming them to a juvenile state (55). Thus, the study of iPSC-derived neurons is limited to young cells, which is suboptimal since age is a key determinant of HD pathology (2, 3). As a consequence, most HD-iPSCs studies with well documented phenotypes are of limited utility (51–54). As an alternative to iPSCs, we and others have recently developed direct lineage reprogramming (25, 56). By overexpressing and knocking-down key transcription factors it is possible to reprogram human fibroblasts directly into neurons, without going through a juvenile state. This approach allows for the generation of patient-derived neurons that retain age-associated epigenetic marks (21–23).

In this study we have used direct reprogramming of patient-derived fibroblasts into iNs to study disease mechanisms in HD. As mentioned above, this approach has many advantages for the study of neurodegenerative disorders as it allows for easy access of patient-derived neurons that retains ageing-related epigenetic marks of the donor and which can be studied in detail using a wide panel of molecular and biochemical technologies. Using this approach, we were able to detect clear disease-related phenotypes when studying iNs from individuals with CAG repeats in the range normally seen in clinic in patients (4).

In HD-iNs we found that there is a post transcriptionally altered cell-type specific autophagy impairment and sought to understand the mechanism by which this comes about. Several studies have demonstrated that the presence of mHTT interrupts autophagy, contributing to the impaired clearance of aggregated proteins (6–10, 57). In various models of HD, different kinds of impairments in autophagy have been described including an increased number of autophagosomes (which sometimes appear empty), disrupted vesicle trafficking and impaired autophagosome-lysosome fusion and dynamics (7, 8, 10, 58). It is also not clear if impaired autophagy directly contributes to the buildup of protein aggregates or if the aggregates themselves influence the activity of autophagy (7-10, 42, 59-63). It has been speculated that defects in the autophagic machinery can lead to a negative feedback loop, whereby mHTT aggregation leads to a further dysregulation of autophagy causing increased mHTT accumulation and neurotoxicity (7, 63–65). Thus, while there are numerous experimental reports on autophagy impairments in HD, it remains unclear which of these are specific to the model system and which are relevant to the actual disease (7-10, 42, 59-63). This is important given its therapeutic implications and the fact that trials are now starting to appear in the clinic looking at autophagy enhancing agents. In HD-iNs, we found a subcellular, neurite specific autophagy impairment, with an accumulation of LAMP1-positive late autophagic structures. We also show that this is a consequence of an impaired transport of these structures to the cell soma where they should be degraded. This finding provides an answer as to why neurons are particularly vulnerable in HD and represent a novel therapeutic target – restoration of autophagolysosome transfer to the cell soma.

The underlying molecular mechanism for the autophagy impairment in HD-iNs appear to be linked to the AMPK pathway, since several factors in this pathway were dysregulated. AMPK is a key energy sensor that promotes catabolic pathways while shutting down ATP consuming processes required for cell growth (66–68). AMPK inhibits cell growth by inhibiting mTORC signaling and protein synthesis downstream of mTORC1. Energy impairments such as decreased mitochondrial biogenesis and trafficking, oxidative stress, increased apoptosis, and ATP deficit all have been implicated in HD pathogenesis (49). Neurons are energetically demanding cells and thus highly vulnerable to abnormalities in cellular respiration. Our findings point towards boosting autophagy by specifically targeting the AMPK pathway. In line with this, we and others have also shown that BECN1 overexpression can rescue some aspects of HD pathology in various models (6, 7, 33, 34). Moreover, genetic and pharmacological activation of AMPK has been shown to protect dysfunctional and vulnerable neurons in HD in nematode, cellular and mouse models (69, 70). An impairment of autophagy in neurons will have multiple pathological consequences (13, 14). Autophagy is implicated in neurogenesis, synaptogenesis, the control of post-transcriptional networks and protein aggregation (6, 71–73). Thus, impairment of autophagy could underlie many of the early cellular disease phenotypes observed in HD (74, 75). As such, the development of specific autophagy-boosting therapies is promising as they have the potential to directly restore other dysfunctional intracellular processes.

Since HD-iNs retain ageing epigenetics characteristics, our results indicate that autophagy impairments in HD may be a combination of age-related epigenetic alterations and m*HTT*-mediated post transcriptional processes. Exactly how the presence of a m*HTT*-allele results in a reduction in the transport of autophagolysosomes from neurites remains unknown, but a combination of an age-related alteration in autophagy-control together with a direct m*HTT*-mediated protein-protein interaction appears the most likely scenario. For example, mHTT has previously been found to directly interact and destabilize BECN1, which is in line with the reduction of BECN1 protein that we found in HD-iNs (6, 7, 33, 34, 57, 76). How ageing and the epigenetic alterations influences the disease pathology and autophagy impairments is currently unknown but will be interesting to investigate in order to find mechanistic links between these phenomena.

Our study also has direct implications for the development of therapies working on m*HTT*-silencing. Such therapies are considered a very promising possibility to successfully treat HD-patients and clinical trials are already underway (17, 77, 78). Our results suggest that the development of allele-specific silencing of m*HTT* may be key to the success of such therapies given that wt*HTT* is directly involved in the control of cellular pathways controlling protein degradation. Thus, while the silencing of m*HTT* will certainly have beneficial consequences, as demonstrated in our study by efficiently lowering LAMP1 in the neurites, the silencing of wt*HTT* will also come with loss-of-function consequences on similar cellular pathways.

In summary, we have developed a novel cell-based model of HD that allows for the study of aged patient-derived neurons. We found that HD-iNs display distinct autophagy alterations, characterized by a blockage in autophagolysosome transfer and degradation. Our results thus identify a novel therapeutic target through autophagy while also helps to advocate for the development of allele specific silencing-based HD therapies.

## Methods

### Human Tissue

Post-mortem human brain tissue was obtained from the Cambridge Brain Bank (Cambridge, UK) and used under local ethics approval (REC 01/177). Severity of HD was graded by a certified pathologist according to the Vonsattel grading system (79) (Table 2).

### Cell Culture

Adult dermal fibroblasts were obtained from the Huntington’s disease clinic at the John van Geest Centre for Brain Repair (Cambridge, UK) and used under local ethical approval (REC 09/H0311/88). The cells were obtained from 10 HD and 10 healthy individuals (Table 1), for more information on the biopsy sampling see (21). CAG repeat length was defined for both alleles using Sanger Sequencing (Laragen Sanger Sequencing Services). The fibroblasts were kept in DMEM Glutamax medium (Gibco) supplemented with 10% FBS (Gibco) and 1% penicillin/streptomycin (Gibco) and passaged when they reached 80–90% confluency using a previously described procedure (21).

### Lentiviral production

Third-generation lentiviral vectors were produced as previously described (6).

For iN conversion LV.U6.shREST1.U6.shREST2.hPGK.BRN2.hPGK.Ascl1.WPRE transfer vector was used. This previously published and available construct from the plasmid repository contains the transcription factors *ASCL1* and *BRN2* with two short hairpin RNAs (shRNA) targeting *REST* (21). The lentiviral vector also contains non-regulated ubiquitous phosphoglycerate kinase (*PGK*) promoters and a Woodchuck Hepatitis Virus (WHP) Posttranscriptional Regulatory Element (*WPRE*). Four additional viral vector plasmids were used: pLV.hU6-sgLacZ-hUbC-dCas9-KRAB-T2a-GFP (*LacZ*), pLV.hU6-sg1HTT-hUbC-dCas9-KRAB-T2a-GFP (*g1HTT*), pLV.hU6-sg2HTT-hUbC-dCas9-KRAB-T2a-GFP (*g2HTT*) and pLV.hU6-sg3HTT-hUbC-dCas9-KRAB-T2a-GFP (*g3HTT*). Vectors are specified in the CRISPRi method section later on.

Virus titration was performed, and the titer was determined with qRT-PCR as previously described (6). The virus titers ranged between 2.33E+08 and 9.3E+09. A MOI of 1-20 was used from different lentiviral vectors as specified for each case.

### Neural conversion

Prior to the start of conversion, Nunc Delta surface treated plates (Thermo Scientific) were coated as previously described (24). Fibroblasts were plated at a density of 50,000 cells per Nunc 24-well (approximately 26,000 cells/cm^2^) in fibroblast medium one day prior to the start of conversion. On the following day (day 0), the fibroblasts were transduced with the all-in-one lentiviral vector at MOI 20. The conversion was performed as previously described until the cells were harvested for experiments on day 25, 28 or 50 of conversion as described in the sections below (21).

### CRISPRi

In order to silence the transcription of *HTT* we used the catalytically inactive Cas9 (deadCas9) fused to the transcriptional repressor KRAB (80, 81). Single guide sequences were designed to recognize DNA regions just down-stream of the *HTT* transcription start site (TSS at 3074690 using reference sequence NC_000004.12) according to https://portals.broadinstitute.org/gpp/public/analysis-tools/sgrna-design-crisprai?mechanism=CRISPRi algoritms.

The guides were inserted into a deadCas9-KRAB-T2A-GFP lentiviral backbone containing both the guide RNA under the U6 promoter and dead-Cas9-KRAB and GFP under the Ubiquitin C promoter (pLV hU6-sgRNA hUbC-dCas9-KRAB-T2a-GFP was a gift from Charles Gersbach (Addgene plasmid #71237; http://n2t.net/addgene:71237; RRID:Addgene_71237)). The guides were inserted into the backbone using annealed oligos and the BsmBI cloning site. Lentiviruses were produced as described above yielding titers between 4.9E+08 and 9.3E+09. Three guides were designed and tested in HEK293T and iPS cells. HEK293T cells were cultured similarly to the fibroblasts cells as described above. iPS cells (RBRC-HPS0328, 606A1 from RIKEN) were cultured as previously described (82). HEK293T and iPS cells were transduced with different gRNAs targeting *LacZ* or *HTT*. After 4 days of transduction, cells were passaged and seven days post infection GFP^+^PI^-^ cells were purified by FACS. Silencing efficiency was tested using quantitative real-time PCR and two gRNAs were chosen for further analysis.

Guide 2 and Guide 3 were chosen for further validation with “cut-sites” at 25 bp and 65 bp downstream of the TSS, respectively. Control LacZ virus with a gRNA sequence not present in the human genome was also produced and used in all experiments. All lentiviral vectors were used with a MOI of 20. Cells were FACS sorted one week after transduction and silencing efficiency was validated using standard quantitative real-time reverse transcriptase PCR techniques as described below.

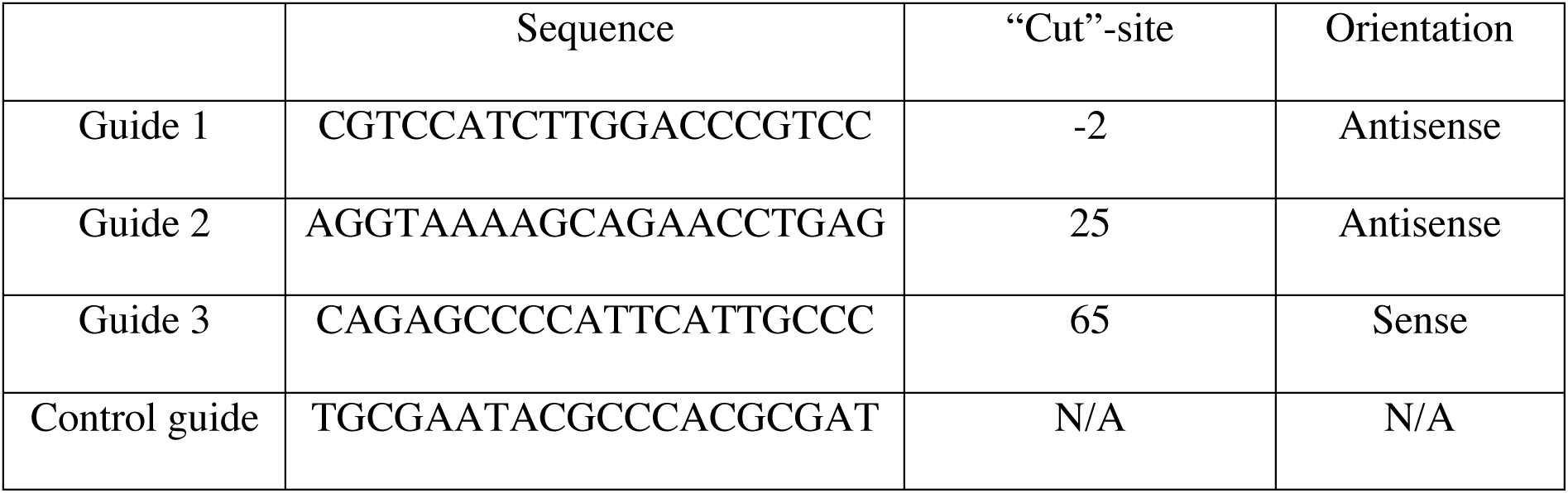

### Autophagy Treatments

Cells were treated with factors regulating autophagy as follows. The cell medium was aspirated from the wells and fresh medium with one of the factors (Bafilomycin, 200 nM, Merck Millipore; Rapamycin, 20 nM, Sigma-Aldrich; Wortmannin, 100 nM, Sigma-Aldrich) was added to the well followed by fixation for ICC after four hours. Non-treated wells received fresh media with DMSO in equivalent amount to that used in treated cells.

Cells were starved by replacing the media with Hank’s Balanced Salt Solution (Thermo Fisher, 14025092) for two hours before fixation.

### Immunostaining

Immunocytochemistry to stain iNs was performed as previously described (21). Briefly, the cells were fixed with 4% paraformaldehyde for 10-15 minutes. Following fixation, the paraformaldehyde was aspirated, and the cells were washed carefully twice with DPBS. Thereafter, the cells were permeabilized in 0.1 M PBS with 0.1% Triton X-100 for 10 min and then blocked for a minimum of 30 min in a blocking solution of 0.1 M PBS and 5% normal donkey serum. The primary antibodies were diluted in blocking solution and incubated overnight at 4 °C (Table 3). The cells were washed twice with DPBS and the secondary antibody conjugated to a fluorophore (Table 3) diluted in blocking solution was added and incubated for 2 hours at room temperature. Following incubation with the secondary antibodies, DAPI was applied for 15 minutes and the cells were washed once with DPBS. Finally, high-content automated microscopy analysis was performed either using the Cellomics Array Scanner (VT1 HCS Reader, Thermo Fischer) and a Leica inverted fluorescent microscope (model DMI6000 B) or a Leica TCS SP8 confocal laser scanning microscope.

**Table 3.** List of antibodies used for ICC and IHC.

| Antibody | Conc. | Species | Company | Cat# | RRID |
| --- | --- | --- | --- | --- | --- |
| **MAP2** | 1:500 | Rabbit | Millipore | Ab5622 | AB_91939 |
| **MAP2** | 1:10,000 | Chicken | Abcam | 5392 | AB_2138153 |
| **P62** | 1:500 | Rabbit | Abcam | Ab91526 | AB_2050336 |
| **LC3B** | 1:500 | Rabbit | Sigma | L7543 | AB_796155 |
| **LAMP1**<br>**(clone H4A3)** | 1:200 | Mouse | DSHB | N/A | AB_2296838 |
| **TAU (clone HT7)** | 1:500 | Mouse | Thermo Scientific | MN1000 | AB_2314654 |
| **TAU** | 1:1,000 | Rabbit | Dako | A0024 | AB_10013724 |
| **Neurofilament**<br>**Protein antibody** | 1:200 | Mouse | Agilent | M0762 | AB_2314899 |
| **Alexa Fluor®**<br>**488 AffiniPure**<br>**Donkey IgG** | 1:200 | Rabbit | Jackson | 711-545-152 | AB_2313584 |
| **Alexa Fluor®**<br>**488 AffiniPure**<br>**Donkey IgG** | 1:200 | Mouse | Jackson | 715-545-150 | AB_2340846 |
| **Cy™3**<br>**AffiniPure**<br>**Donkey IgY** | 1:200 | Chicken | Jackson | 703-165-155 | AB_2340363 |
| **Cy™3**<br>**AffiniPure**<br>**Donkey IgG** | 1:200 | Rabbit | Jackson | 711-165-152 | AB_2307443 |
| **Cy™3**<br>**AffiniPure**<br>**Donkey IgG** | 1:200 | Mouse | Jackson | 715-165-151 | AB_2315777 |
| **Alexa Fluor®**<br>**647 AffiniPure**<br>**Donkey IgG** | 1:200 | Mouse | Jackson | 715-605-150 | AB_2340862 |

Immunohistochemistry staining was performed as described before (6). 10 μm thick paraffin-embedded striatal sections were taken from 3 differently graded HD patients and healthy age-matched control brains. Sections were stained using the antibodies listed in Table 3, using mouse anti-Neurofilament and rabbit anti-p62. Briefly: sections were surrounded with Dakopen and dried for 10 minutes at 65 °C. Afterwards sections were further incubated first with xylene and then with different concentrations of ethanol (99.5 %, 95 %, 70 %), MilliQ water and last in TN buffer (1 M TRIS-HCl, 1.5 M NaCl, MilliQ water) before 20 minutes of boiling in a pH = 9 TRIS/ EDTA solution. After cooling, the sections were again twice incubated with TN buffer and then with TN + 5 % serum in RT. The sections were incubated at RT with the primary antibody diluted in TNT + 5 % serum (TN + 10 % Tween20). Secondary antibodies were diluted in TNT + 5 % serum and kept in the dark for 2 hours after washing with TN and TNT. Lastly, sections were washed, and cover slipped with PVDA-DABCO with DAPI. All fluorescent images were taken using a Leica TCS SP8 confocal laser scanning microscope.

### High-content automated microscopy

The Cellomics Array Scan (VT1 HCS Reader, Thermo Fischer) was used for high-content automated microscopy.

To quantify the number of DAPI^+^, MAP2^+^, and TAU^+^ cells and define neuronal purity and conversion efficiency “Target Activation” (TA) was used. Using this method, we obtained objective, unbiased measurements of the iN cultures. The TA program was used to acquire images of 100-289 fields using a 10x objective of each well to define cell number, neural purity and conversion efficiency. Wells with <50 valid fields were excluded from further analysis. The program defined DAPI^+^ cells based on intensity and nuclei and then measured fluorescent intensity on a cell-by-cell bases to identify MAP2^+^ and TAU^+^ cells. Border objects were excluded from the analysis. The neuronal purity was quantified as the fraction of MAP2^+^ or TAU^+^ cells of the total DAPI^+^ cells at the time of analysis. The conversion efficiency was determined as the number of MAP2^+^/TAU^+^ cells over the number of fibroblasts plated at the start of conversion.

The program “Neuronal Profiling” (NP) was used at a 10x objective. NP analysis was performed by re-analyzing images taken for the TA analysis to quantify the neuronal morphology of the MAP2^+^ and TAU^+^ cells. Control and HD-iNs were imaged with a 10x objective after 50 days of conversion. The NP program was used to acquire images of 100-289 fields at 10x magnification of each well to define neuronal morphology. Wells with <50 valid fields were excluded from further analysis. First valid nuclei were defined by DAPI staining based on intensity and area. Border objects were excluded from the analysis. Average cell body area, average neurite area, average number of neurites, average neurite length, average neurite width and average number of branchpoints per cell was defined by the NP program based on MAP2 or TAU neuronal staining. Border objects were excluded from further analysis.

Neuronal Profiling with spot detection (using a 20x objective) was used to determine average LC3B, LAMP1 and p62 dot number and size per cell within MAP2^+^ or TAU^+^ cell bodies and neurites. LAMP1 and LC3B colocalization was also analyzed by defining the overlapping area as a percentage between the two markers. First valid nuclei were defined by DAPI staining based on intensity and area. Border objects were excluded from further analysis. Next, cell bodies and neurites were defined based on intensity and area of MAP2^+^ or TAU^+^ as the region of interest. Border objects were excluded from further analysis here also. Autophagy markers were analyzed and defined by intensity and area within the valid neuronal cells. In every case, 150-250 fields were analyzed and wells <50 valid fields were excluded from further analysis.

### FACS

To increase the purity of converted control and HD-iNs for RNA-sequencing, the cells were harvested and sorted by FACS as previously described (24). To this end, the cells were dissociated with StemPro accutase by incubation at 37 °C for approximately 10-15 min. Following detachment, the cells were washed off and collected in FACS buffer (HBSS with 1% BSA and 0.05% DNAse I (Sigma)) and centrifuged at 400 x g for 5 min. The supernatant was aspirated, and the cells were once more washed in FACS buffer, centrifuged, and resuspended in 50 *µ*l FACS buffer. An allophycocyanin conjugated antibody against human NCAM (1:10, anti-mouse hNCAM, Biosciences, #555515, clone B159) was added to the samples and incubated for 15 minutes on the bench. The antibody was washed twice with FACS buffer and centrifuged again with the same settings. After the final dilution, 1:1,000 propidium iodide (PI, Sigma) was added to label dying cells. The cells were sorted with a FACSAria III through a 100 µm nozzle, using a 1.5 filter and area scaling of 0.35. Gates were set up to obtain small NCAM^+^PI^-^ cells using fluorophore specific gates and the forward and side scatter to select the smaller cell population. Re-analysis was also performed for each sorted sample and a purity >95% was set as a cutoff. In each case 10,000 NCAM^+^PI^-^ single cells were sorted at 10 °C and the samples were then kept on ice for further processing. Sorted cells were centrifuged at 400 x g for 5 minutes and after the removal of the supernatant, frozen on dry ice and stored for RNA-sequencing experiments.

To purify successfully transduced GFP^+^ HEK293T, iPSCs or fibroblasts, these cells were also harvested and sorted by FACS. Untransduced and transduced cells after one week of lentiviral transduction were dissociated with 0.05 % trypsin (Sigma) for 5 minutes at 37 °C. Following detachment, the cells were washed off and collected in FACS buffer and centrifuged at 400 x g for 5 minutes. Supernatant was removed and cells were washed with a FACS buffer again. After washing cells were filtered through a 60 µm sterile nylon filter to remove possible cell aggregates and collected in 500 µl FACS solution. Before sorting, cells were stained with PI. GFP^+^PI^-^ cells were sorted into fresh DMEM medium for further analysis. In all cases untransduced cells were also FACS sorted. Re-analysis was also performed for each sorted sample and a purity >95 % was set as a cutoff.

### Quantitative Real-time PCR

To measure the expression level of *HTT* RNA in fibroblasts and iNs from healthy controls and HD patients, we did qRT-PCR analysis. Total RNA was first extracted according to the supplier’s recommendations using the mini or micro RNeasy kit (Qiagen). cDNA was generated using the Maxima First Strand cDNA Synthesis Kit. All primers were used together with LightCycler 480 SYBR Green I Master (Roche). Three reference genes were used for each qRT-PCR analysis (ACTB, GAPDH and HPRT). Sequences were:

*ACTB*

fw: CCTTGCACATGCCGG

rev: GCACAGAGCCTCGCC

*GAPDH*

fw: TTGAGGTCAATGAAG

rev: GAAGGTGAAGGTCGG

*HPRT*

fw: ACCCTTTCCAAATCCTCAGC

rev: GTTATGGCGACCCGCAG

*HTT* expression levels were tested using two alternative primer pairs. Sequences were:

*HTT* – pp1

fw: TCAGCTACCAAGAAAGACCGT

rev: TTCCATAGCGATGCCCAGAA

*HTT* – pp2

fw: TCAGAAATGCAGGCCTTACCT

rev: CCTGGACTGATTCTTCGGGT

In all cases data were quantified using the ΔΔCt method (REF).

### Western Blot

Fibroblasts (200,000 cells/ sample) and iNs (converted in T25 flasks starting from 250,000 fibroblasts/ sample or converted in Nunc Delta treated 6-well plates starting from 200,000 plated fibroblasts/ sample) were harvested as follows: the cell medium was removed and the cells were lysed in RIPA buffer (Sigma) with 4% protease inhibitor cocktail (PIC, cOmplete). The lysed cells were collected in a microcentrifuge tube and incubated on ice for a minimum of 30 minutes, followed by centrifugation at 10,000 g for 10 minutes in 4 °C to pellet cellular debris. Following centrifugation, the supernatant was transferred to new vials. The protein content was quantified with Bradford DC^TM^ Protein Assay (Biorad) and 10-15 µg protein of each sample was used for loading the gel. Gel electrophoresis and blotting was performed as previously described (7). Both the primary and secondary antibodies (Table 4) were diluted in milk blocking solution. The blots were incubated in Immobilon Western Chemiluminescent HRP Substrate (Millipore) for 5 minutes to enhance the signal for visualization using the ChemiDoc^TM^ MP Imaging System.

**Table 4.** List of antibodies used for WB.

| Antibody | Conc. | Species | Company | Cat# | RRID |
| --- | --- | --- | --- | --- | --- |
| **BECN1** | 1:500 | Rabbit | Santa Cruz | sc-11427 | AB_2064465 |
| **P62** | 1:5,000 | Mouse | Abcam | ab56416 | AB_945626 |
| **LC3B** | 1:5,000 | Rabbit | Sigma | L7543 | AB_796155 |
| **LAMP1**<br>**(clone H4A3)** | 1:1,000 | Mouse | DSHB | N/A | AB_2296838 |
| **Hrp Conjugated**<br>**antibody** | 1:2,500 | Rabbit | Sigma-Aldrich | NA9340 | AB_772191 |
| **Hrp Conjugated**<br>**antibody** | 1:5,000 | Mouse | Santa Cruz | Sc-2005 | AB_631736 |
| **β-actin** | 1:100,000 | Mouse | Sigma-Aldrich | A3854 | AB_262011 |

To determine HTT protein expression in fibroblasts and iNs protein concentration was determined using Bradford Protein Assay (Bio-Rad Laboratories, Mississauga, ON, Canada). HTT immunoblotting was performed as previously described (83). Briefly, proteins were loaded on a Tris-Acetate gradient gel (3-10% 37.5 :1 acrylamide/Bis-acrylamide, BioShop, Burlington, ON, Canada), migrated at 100V and transferred in Bicine/Bis-Tris transfer buffer overnight at 25V and 4°C, followed by 1 hour at 90V. Membranes were blocked with 5% milk and 1% bovine serum albumin in Tris-buffered saline with 0.1% Tween 20. Membranes were than incubated with primary antibodies raised against poly-glutamine repeats (5TF1-1C2, EMD millipore) or HTT (1HU-4C8 and mEM48, both from EMD Millipore; CH00146, CHDI – Corriell Institute). Detection was achieved using appropriate horseradish peroxidase-labeled secondary antibodies and Immobilon Western chemiluminescent HRP substrate (Millipore Sigma). Band intensity was determined with ImageJ 2.0.0-rc-69/1.52p software (http://imagej.nih.gov/ij) and corrected to the total amount of protein per lane.

### RNA preparation and sequencing

Samples C1-C7 and HD1-2, HD5-7, and HD9-10 were used for RNA-sequencing. Total RNA was extracted from 10,000 cells/ sample with the RNeasy micro kit (Qiagen) according to the manufacturer’s protocol. A quality control of the samples was made with the Bioanalyzer RNA pico kit. cDNA was synthesized with the SMART-Seq® v4 Ultra® Low Input RNA Kit for Sequencing (Takara/Clontech) and assessed with the Bioanalyzer high sensitivity DNA kit, followed by library preparation using Nextera XT (Illumina). The quality and concentration of the libraries was assessed with the Bioanalyzer high sensitivity DNA kit and Qubit dsDNA BR DNA assay kit, respectively. Paired-end sequencing of 2 x 150 base pairs (300 cycles) was done with a NextSeq 500/550 High Output v2.5 kit 400 million reads (Illumina) on a NextSeq 500 sequencer (Illumina).

### RNA sequencing analysis

Fibroblast and iN samples were sequenced as specified above for further analysis. Raw base counts were demultiplexed and converted to sample-specific fastq format files using the bcl2fastq program (Illumina) with default parameters. The quality of the reads was assessed using FastQC (https://www.bioinformatics.babraham.ac.uk/projects/fastqc/) and MultiQC (https://multiqc.info/), after which reads were mapped to the human genome (GRCh38) using the STAR mapping algorithm with default parameters (84). Following mapping, mRNA expression was quantified using FeatureCounts. Only reads mapping to genetic elements annotated as exons were quantified, and only the primary alignments were included (85). The GTF annotation file used for the quantification was downloaded from Gencode version 30 (https://www.gencodegenes.org/human/). We performed median of ratios normalization with DESeq2 (86) to account for differences in sequencing depth and RNA composition. Gene ontology overrepresentation tests were performed using PANTHER database (version 14). The GO analysis between iNs and fibroblasts was performed using the up and downregulated genes found to be significantly different using DESeq2. Genes with basemean more than 10 were used as the background set for the overrepresentation test, and only significant terms are shown (padj <0.05, log2FC >1).

Using the normalized reads (mean of ratios calculated with DESeq2), we tested for difference in expression between HD and ctrl-iNs using unpaired t-test. We defined significantly different genes those with p-value<0.05. Code for tests and visualization is available at Github (https://github.com/ra7555ga-s/iN_HD) (86). Gene ontology overrepresentation tests comparing HD and ctrl-iNs using the RNA sequencing data were performed using PANTHER (version 14) using Fisher’s exact test and Benjamini-Hochberg correction to calculate false discovery rates (87). All genes with some expression in any of the conditions were used as background sets for these tests.

### Shotgun proteomic analysis

Samples C1-C7 and HD1-2, HD5-7, and HD9-10 were used for mass spectrometry (MS) analysis. Fibroblasts (500,000 cells) and iNs converted in T75 flasks (600,000 fibroblasts plated for conversion per sample) were dissociated as previously described and prepared for quantitative proteomic analysis as follows. The cells were carefully washed off and collected in a tube with either trypsin or accutase and spun at 400 x g for 5 minutes. The supernatant was discarded, and the pellets were washed three times with DPBS. After the final wash, the supernatant was aspirated, and the pellets were frozen on dry ice and stored at −80 °C until use. The cell pellets were resuspended in 200 µL lysis buffer (50 mM DTT, 2 %SDS, 100 mM Tris pH = 8.6), rested for 1 min on ice and sonicated (20 cycles: 15 seconds on/off; Bioruptor plus model UCD-300, Diagenode). Reduction and alkylation of disulfide bridges was performed by incubating the samples at 95 °C for 5 minutes, followed the addition of iodoacetamide to a final concentration of 100 mM and incubation for 20 minutes at room temperature in the dark.

Samples were processed using S-Trap Mini Spin Columns (ProtiFi, USA) according to the manufacturer’s instructions. Briefly, samples were acidified by adding phosphoric acid to a final concentration of 1.2%, 7 volumes of binding buffer (90% MeOH, 100 mM TEAB, pH = 7.1) was added to the samples, which were then transferred to the S-Traps, and spun at 4000 x g for 30 seconds. The trapped proteins were washed three times with the binding buffer. Protein digestion was performed by adding trypsin (Promega Biotech AB) 1:50 (enzyme:protein ratio) in 125 *µ*L of 50 mM TEAB and incubating for 16 hours at 37 °C. Peptides were eluted with 0.2% of aqueous formic acid and 0.2% of formic acid in 50:50 water:acetonitrile. Following speed vacuum concentration peptides were dissolved in 0.1% TFA, quantified with the Pierce Quantitative colorimetric peptide assay (Thermo Fisher Scientific), and 1 µg was injected on the LC-MS/MS system.

Peptides were analyzed in a Dionex Ultimate 3000 RSLCnano UPLC system in line-coupled to a Q-Exactive HF-X mass spectrometer (Thermo Fischer Scientific). Peptides were first trapped on an Acclaim PepMap100 C18 (3 µm, 100 Å, 75 µm i.d. × 2 cm, nanoViper) trap column and separated following a non-linear 120-minute gradient on an EASY-spray RSLC C18 (2 µm, 100 Å, 75 µm i.d. × 25 cm) analytical column. The flow rate was 300 nL/min and the temperature was set to 45 °C. A top 20 data-dependent acquisition (DDA) method was applied, where MS1 scans were acquired with a resolution of 120,000 (@ 200 m/z) using a mass range of 375-1500 m/z, the target AGC value was set to 3E+06, and the maximum injection time (IT) was 100 milliseconds. The 20 most intense peaks were fragmented with a normalized collision energy (NCE) of 28. MS2 scans were acquired at a resolution of 15,000, a target AGC value of 1E+05, and a maximum IT of 50 milliseconds. The ion selection threshold was set to 8E+03 and the dynamic exclusion was 40 seconds, while single charged ions were excluded from the analysis. The precursor isolation window was set to 1.2 Th. Each sample was analyzed in triplicates.

Protein identification and relative label-free quantification was performed by means of the Proteome Discoverer v2.2 (Thermo Fisher Scientific) using SEQUEST HT as the search engine and a human protein database download from UniProt on 2019-01-15. For the search trypsin was selected as the protease, two missed cleavages were allowed, the tolerance was fixed at 10 ppm for MS1 and 0.02 Da for MS2, carbamidomethyl-cysteine was set as static modification while methionine oxidation, phosphorylation on serine, threonine and tyrosine, and protein N-terminal acetylation were selected as dynamic modifications. Peptides and corresponding proteins were identified with 1% of FDR.

Protein quantitative data was processed using Perseus v1.6.5.0. All quantitative values were log2 transformed and standardized by subtracting the median in each sample. The technical replicates were averaged and only those proteins with quantitative values in at least four out of the seven samples, in at least one group were kept for further analysis. The resulting number was 7,001 different proteins. Statistical tests, principal component and 2D functional annotation enrichment analyses were performed in Perseus. The R bioconductor package *limma* was used to fit a linear model and to compute moderated t-statistics (88).

Figures showing scatter plots with mean protein abundance between different cell types (iN and fibroblasts) or conditions (HD and ctrl-iNs) show unpaired t-tests results. The code for these tests and visualization is available on GitHub (https://github.com/ra7555ga-s/iN_HD), as well as the visualization for the functional enrichment analysis (Figure 2e), which was performed with STRING version 11 (89).

### DNA methylation Array and analysis

Samples C1-C3, C5, C6, C9 and HD1-HD8, HD10 were used for DNA methylation analysis. DNA was extracted from ctrl and HD-iNs converted in T25 flasks (200,000 fibroblasts cells plated for conversion per sample) using DNeasy Blood & Tissue Kit (Qiagen) following the manufacturer’s instructions.

Bisulfite conversion was performed using the EZ DNA MethylationTM Kit from Zymo Research. Product No: D5004 with 250 ng of DNA per sample. The bisulfite converted DNA was eluted in 15 μl according to the manufactureŕs protocol, evaporated to a volume of <4 μl, and used for methylation analysis using the Illumina Methylation EPIC array.

For analysis of methylation data, the statistical software R (version 4.0.3) was used. The “minfi” (90) package (version 1.34) was used to determine the quality of methylation experiments and to derive single probe scores per sample after normalization using the preprocessNoob() function (91). All samples were deemed to have acceptable quality based on density plots of beta-values as well as signal intensities for control probes in the red and green channel. Sample age was derived from normalized beta values using the getAgeR function from the cgageR package (https://github.com/metamaden/cgageR).

### Statistical analysis

In iN conversion efficiency and purity analysis each dot represents either one ctrl or a HD adult human fibroblast cell line converted into iNs. Each cell line value is an average from several individual wells specified in each case in the figure legend panels. The well values are generated with high-content automated microscopy analyses. For “Target activation” analysis we scanned at least 100 fields (there are 189 fields in total). We excluded all wells having less than <50 valid fields. MAP2 and TAU purity % was defined for each line by taking the average of:

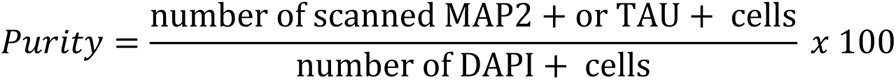

Conversion efficiency % was defined by:

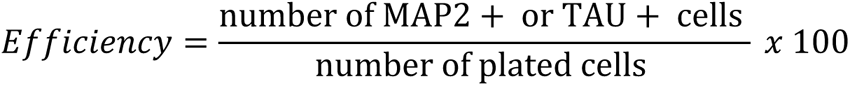

Number of plated cells in a 24-well plate was 50,000 cells.

For all iN morphology analysis each dot represents one control, or one HD adult human fibroblast cell line converted into iNs. Each cell line value is an average from several individual wells specified in each case in the figure legend. The average value for one cell line was defined by: (average value of all wells/ line) / (average value of all control wells performed with identical HCS settings and ICC stainings). All NP values (cell body area, neurite area, count, width, length and branchpoint count) are average relative values per cell.

In all qRT-PCR experiments five control and five HD fibroblast and iN cell lines were analyzed using qRT-PCR by using three different reference genes. In experiments using 293T and iPSCs cell lines we used to test the silencing efficiency of g1HTT, g2HTT and g3HTT using qRT-PCR by using three different housekeeping genes.

In all autophagy measurements each dot represents one control or one HD adult human cell line from the converted iNs. Relative dot number and area values were defined by: (average dot number or area of all wells/ line) / (average dot number or area of all control wells performed with identical HCS settings and ICC staining). Fold changes were defined for each line by first setting every individual non-treated cell line value to 1. Afterwards BAF, W, RAP, ST, g2HTT or g3HTT treated values were defined by:

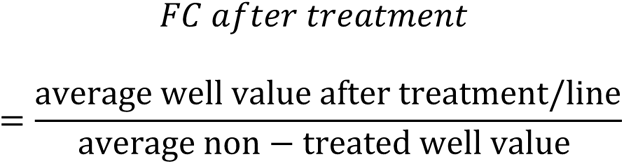

Correlation analysis were tested with Pearson’s correlation coefficient. Correlation between the predicted age (based on Horvath clock) and real age of the donors was tested using Pearson’s correlation coefficient. Two-tailed unpaired T-tests or Paired t-tests were used to test differences between two groups. One-way ANOVA or nonparametric Kruskal-Wallis test was used depending on whether the data obeyed a normal distribution as defined by the D’Agostino-Pearson omnibus normality test to test differences between more than two groups. Two-way ANOVA corrected for multiple comparisons using Sidak statistical hypothesis testing was used when comparing values after various treatments. Multiplicity adjusted p-values were defined for each comparison. Data are presented as min/max box plots or mean and error bars which represent either standard error of the mean (SEM) or standard deviation of the mean (SD), specified in each figure legend.

## Acknowledgements

We are grateful to all members of the Jakobsson lab and Elena Cattaneo for stimulating discussions and useful comments on the manuscript. We also thank U. Jarl, J. Johansson, M. Persson Vejgården and E. Ling for technical assistance. We also thank Dr. Anna Hammarberg for her valuable help with high-content automated microscopy and FACS experiments. Methylation profiling was performed by the SNP&SEQ Technology Platform in Uppsala (www.genotyping.se). The facility is part of the National Genomics Infrastructure supported by the Swedish Research Council for Infrastructures and Science for Life Laboratory, Sweden.

## Author contributions

Conceptualization, K.P. and J.J.; Methodology, K.P., J. D-O., D.A.G., J.G. M.R, B.A.H, K.S., S.L., V.H., J.L., P.A.J, and I.S-A.; Software, R.G., Y.S, D.A.G., P.S., J.G.V and M.R.; Validation, K.P and J.J.; Formal Analysis, K.P, D.A.G., J.G.V, M.R., R.G., Y.S., B.A.H., K.S., J.L., S.L., P.S., V.H., P.A.J., I.S-A., S.S.S. and J.J; Investigation, K.P., J. D-O., D.A.G., S.L., R.G., P.S., V.H., J.G., M.R., Y.S., B.A.H., K.S., J.L., P.A.J, I.S-A., M.P., S.S.S., R.A.B. and J.J.; Resources, K.P., J.G., M.R., P.A.J, Y.S., J.D-O., K.H., R.V., T.S., E.C., M.P., Gy.M-V., S.S.S, R.A.B and J.J; Data Curation, K.P., D.A.G., J.G., M.R., R.G., Y.S., B.A.H, K.S., J.L., S.L., P.S., V.H., P.A.J., I.S-A.; Writing – Original Draft, K.P., R.A.B. and J.J.; Writing – Review & Editing, all authors; Visualization K.P., D.A.G., J.G. and M.E.J.; Supervision, K.P., J. D-O., M.P., S.S.S., Gy. M-V., R.A.B. and J.J.; Funding Acquisition, K.P., R.A.B. and J.J.

## Supplementary Information

**SFigure 1.**
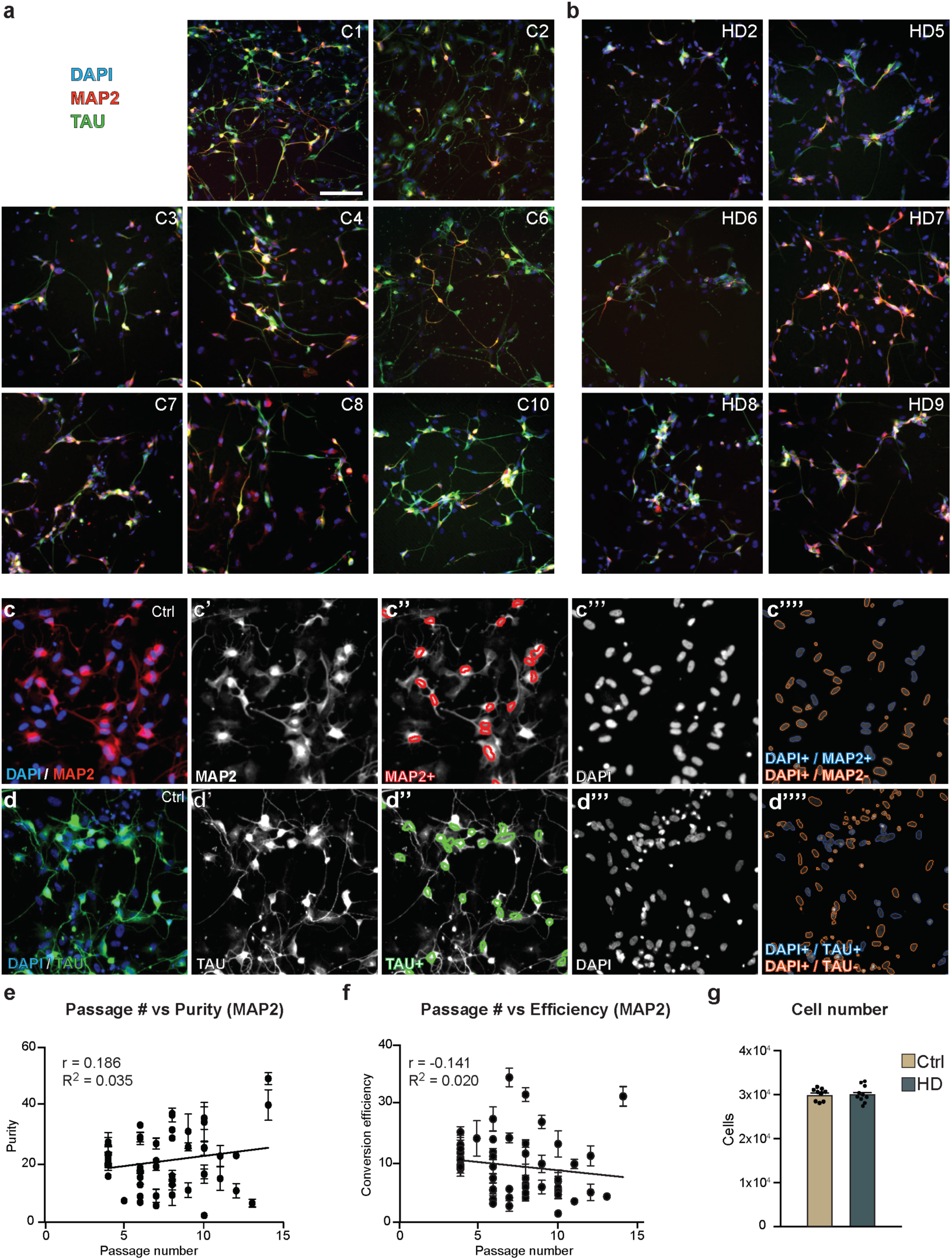
iNs display a neuronal profile. Related to Figure 1. (a-b) Fibroblast-derived iNs from control and HD patients express mature neuronal markers like TAU and MAP2. (c-d) Target activation image analysis by high-content automated microscopy. DAPI^+^ cells are defined by intensity and area criteria. Border objects are excluded. MAP2^+^ and TAU^+^ iNs are defined by average and total intensity measurements on a cell-by-cell basis. On c’’’’and d’’’’ blue circles around the nuclei define the valid MAP2^+^ or TAU^+^ iNs, while orange circled nuclei are not positive for MAP2 or TAU and therefore are not counted as converted neurons. Red circles represent MAP2^+^ iNs in c’’, while TAU^+^ iNs are marked with green circles in d’’. (e-f) There is no correlation between purity or conversion efficiency and passage number (n = 48 replicates; 195 wells analyzed in total). (g) Average DAPI^+^ cell numbers analyzed with high-content automated microscopy. Each dot represents the average value for one control or HD cell line (n = 9 lines for controls, 81 wells analyzed in total; n = 10 lines for HD, 85 wells analyzed in total). (*p<0.05; Two-tailed Pearson’s correlation coefficients were used in e and f. Two-tailed unpaired T-tests were used in g) All data are shown as mean ± SEM. Scale bar is 50 μm.

**SFigure 2.**
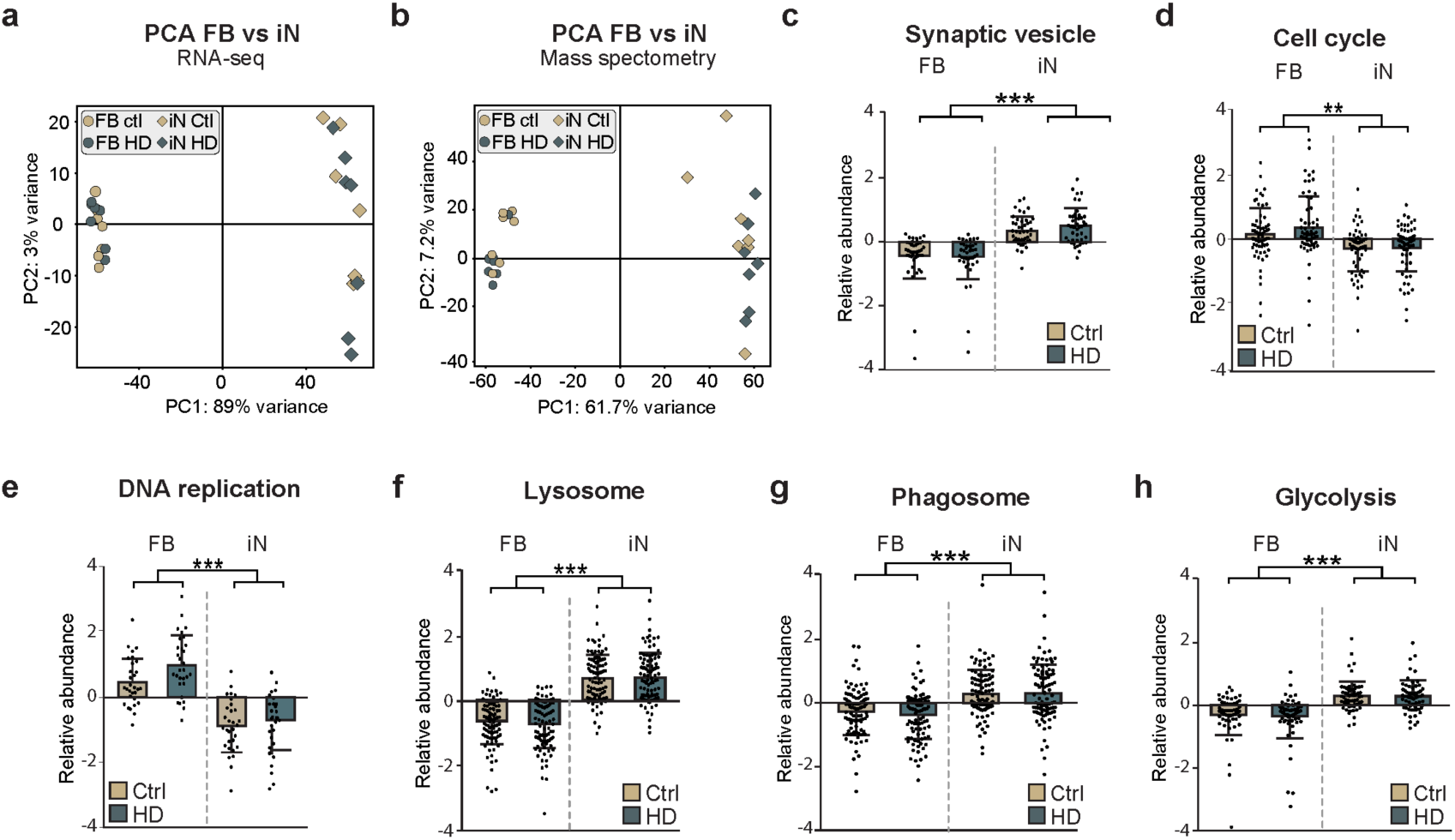
iNs display a neuronal profile at the transcriptome and proteome level. Related to Figure 1. (a) Principal component analysis of transcriptome data of ntop = 500 “top 500 variable genes” showed a major transcriptional difference between fibroblasts and iNs (n = 7 control and 7 HD fibroblast and iN lines). (b) Principal component analysis using the protein abundance profiles of all samples under study. The highest variability explained by the first component separates iNs from fibroblasts. There is more variability among iNs than among fibroblasts (n = 7 control and 7 HD fibroblasts and iN lines). (c-h) Pathways dysregulated between fibroblasts and iNs. Relative abundance profiles of proteins from several pathways (Cell cycle, DNA replication, Glycolysis, Lysosome, Phagosome and Synaptic vesicle) in the four group of samples (n = 7 control and 7 HD fibroblasts and iN lines). (***p<0.001; **<0.01; two-tailed paired T-tests were used in c-h. Data shown as mean ± SD)

**SFigure 3.**
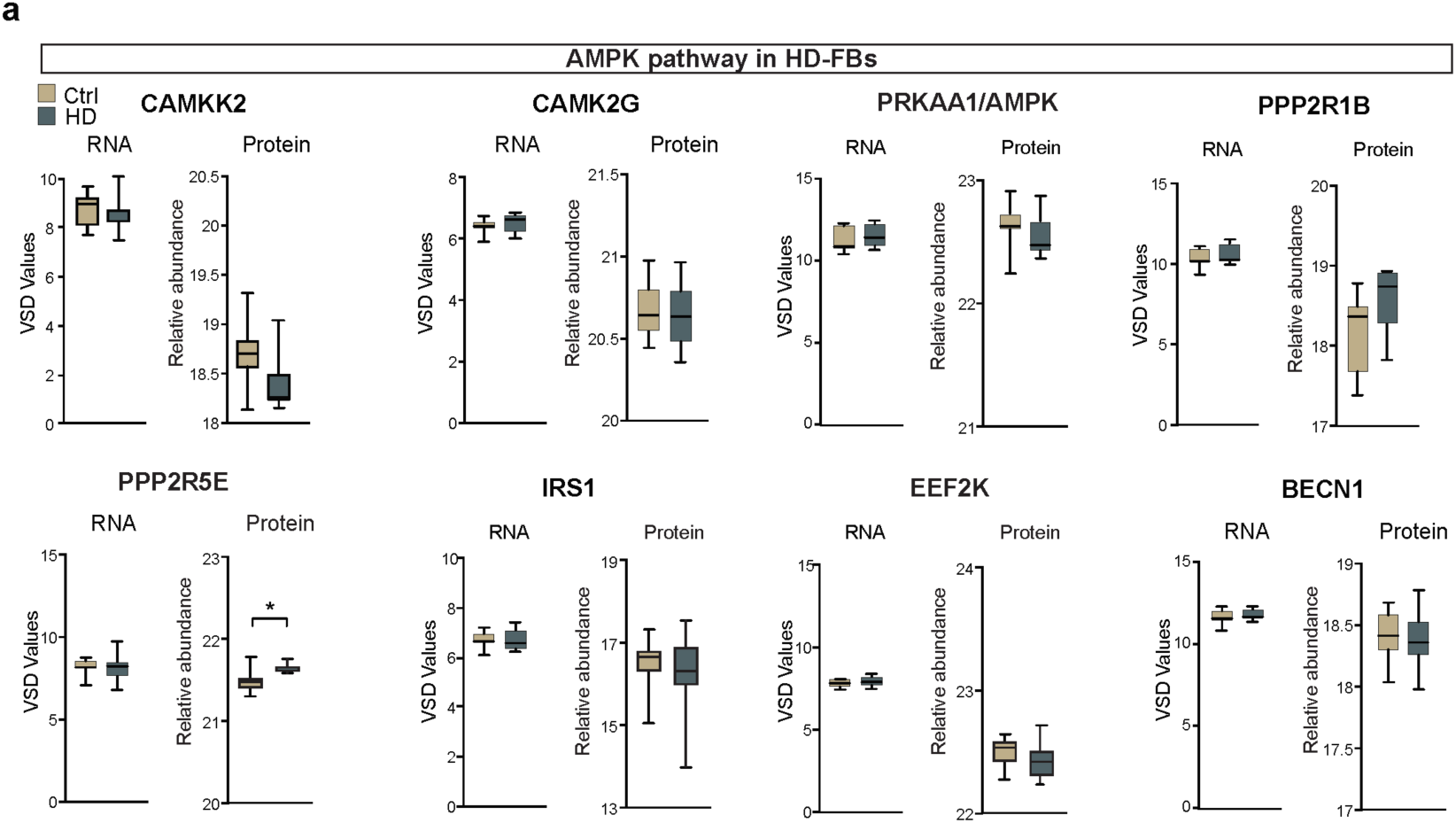
Related to Figure 2. (a) Protein abundance and RNA expression of CAMKK2, CAMK2G, AMPK, PPP2R1B, PPP2R5E, IRS1, EEF2K and BECN1 in control and HD-FBs (n = 7 control and 7 HD fibroblast lines). (*p<0.05; two-tailed unpaired T-tests were used) All data are shown as min/max box plots.

**SFigure 4.**
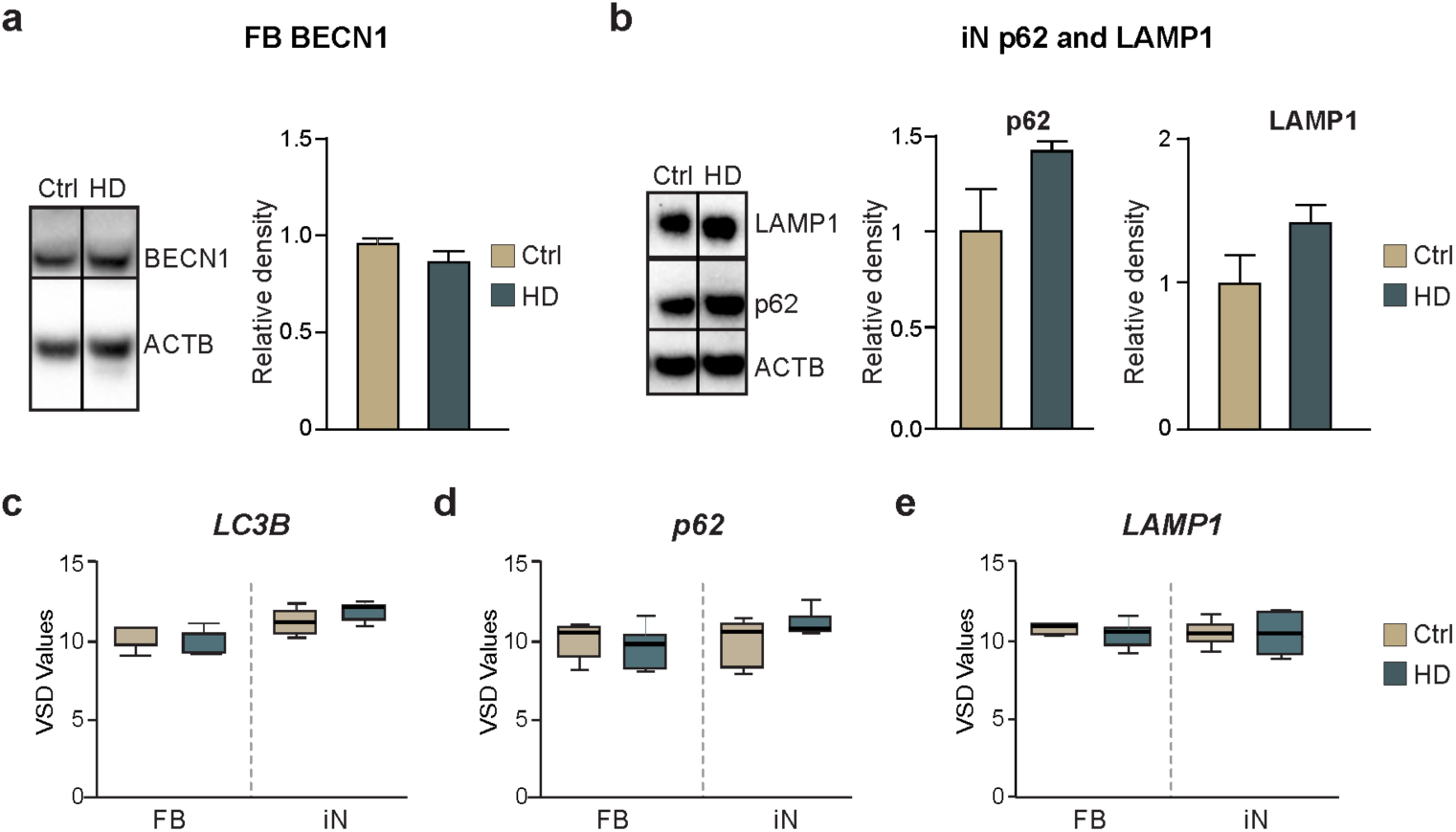
Related to Figure 3. (a) BECN1 expression was not changed in ctrl and HD fibroblasts using WB (n = 10 replicates). (b) p62 and LAMP1 expression was similar between the control and HD-iNs (n = 6 replicates). (c-e) VSD values of *LC3B*, *p62* and *LAMP1* from RNA sequencing of control and HD fibroblasts and iNs (n = 7 control and 7 HD fibroblasts and iN lines). (*p<0.05; two-tailed unpaired T-tests were used). All data are shown as mean ± SEM in a and b. All data are shown as min/max box plots in c. WB values were normalized to non-treated control fibroblasts or iN expression levels and corrected to actin values.

**SFigure 5.**
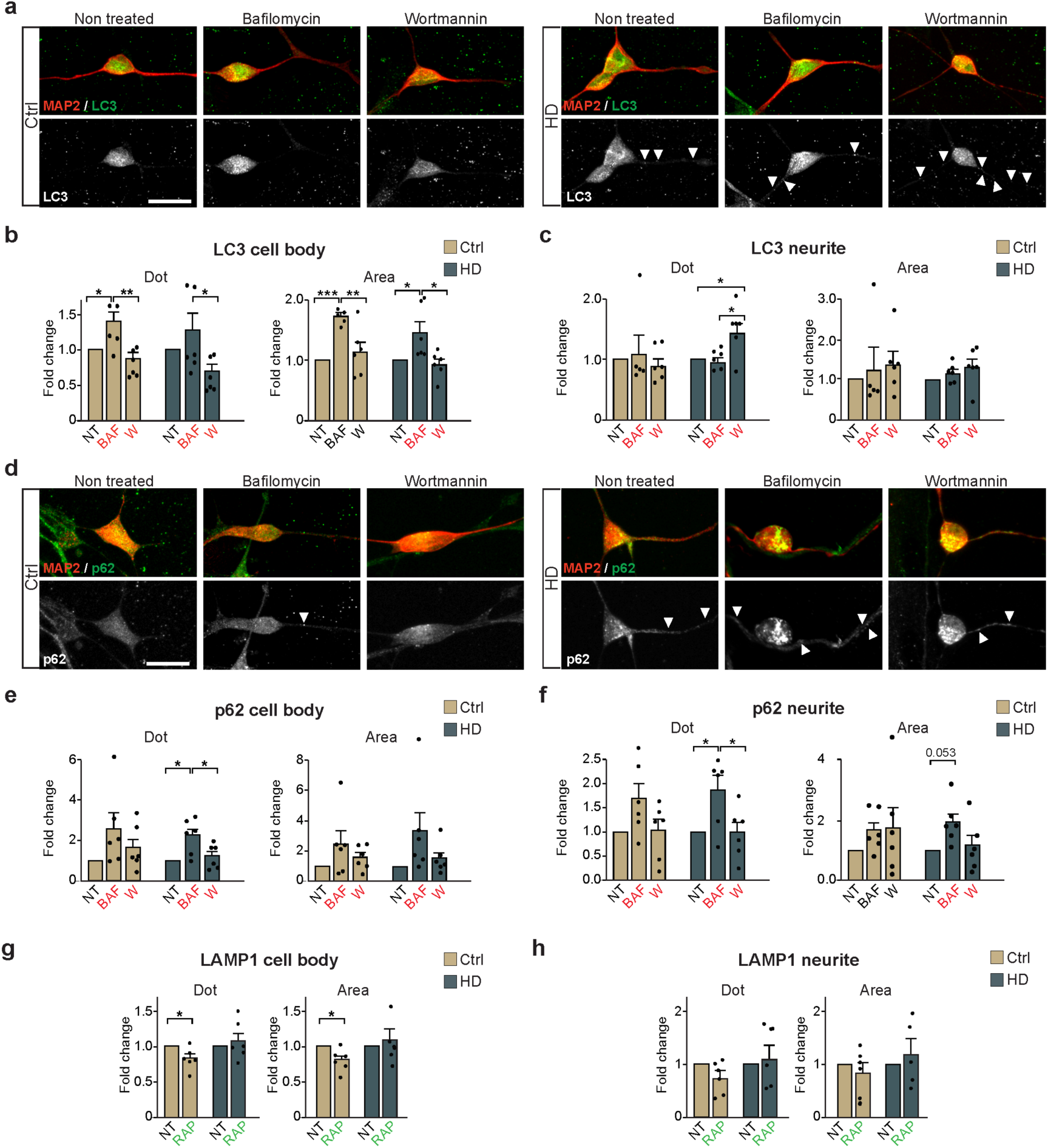
Related to Figure 4. (a-c) Representative images and fold changes of LC3B spot count and area in the cell body and in the neurites in control and HD-iNs after Bafilomycin A1 (Baf) and Wortmannin (W) treatment. Non-treated (-) samples were set to 1 (n = 6 ctrl and HD-iN lines). (d-f) Representative images and fold changes of p62 spot count and area in control and HD-iN cell bodies and neurites after BAF and W treatment (n = 6 ctrl and HD-iN lines). (g-h) Statistical analysis after rapamycin treatment of LAMP1 spot number and size in control and HD-iNs. Non-treated (-) samples were set to 1 (n = 6 ctrl and HD-iN lines). (***p<0.001; **<0.01; *p<0.05; One-way ANOVA or nonparametric Kruskal-Wallis test was used depending on normal distribution defined by D’Agostino-Pearson omnibus normality test for b-c and e-f. Two-tailed paired T-tests were used in g and h). Data are shown as mean ± SEM. Scale bar is 25 μm. FC: Fold-change; -: Non-treated; BAF or B: Bafilomycin A1; W: Wortmannin

**SFigure 6.**
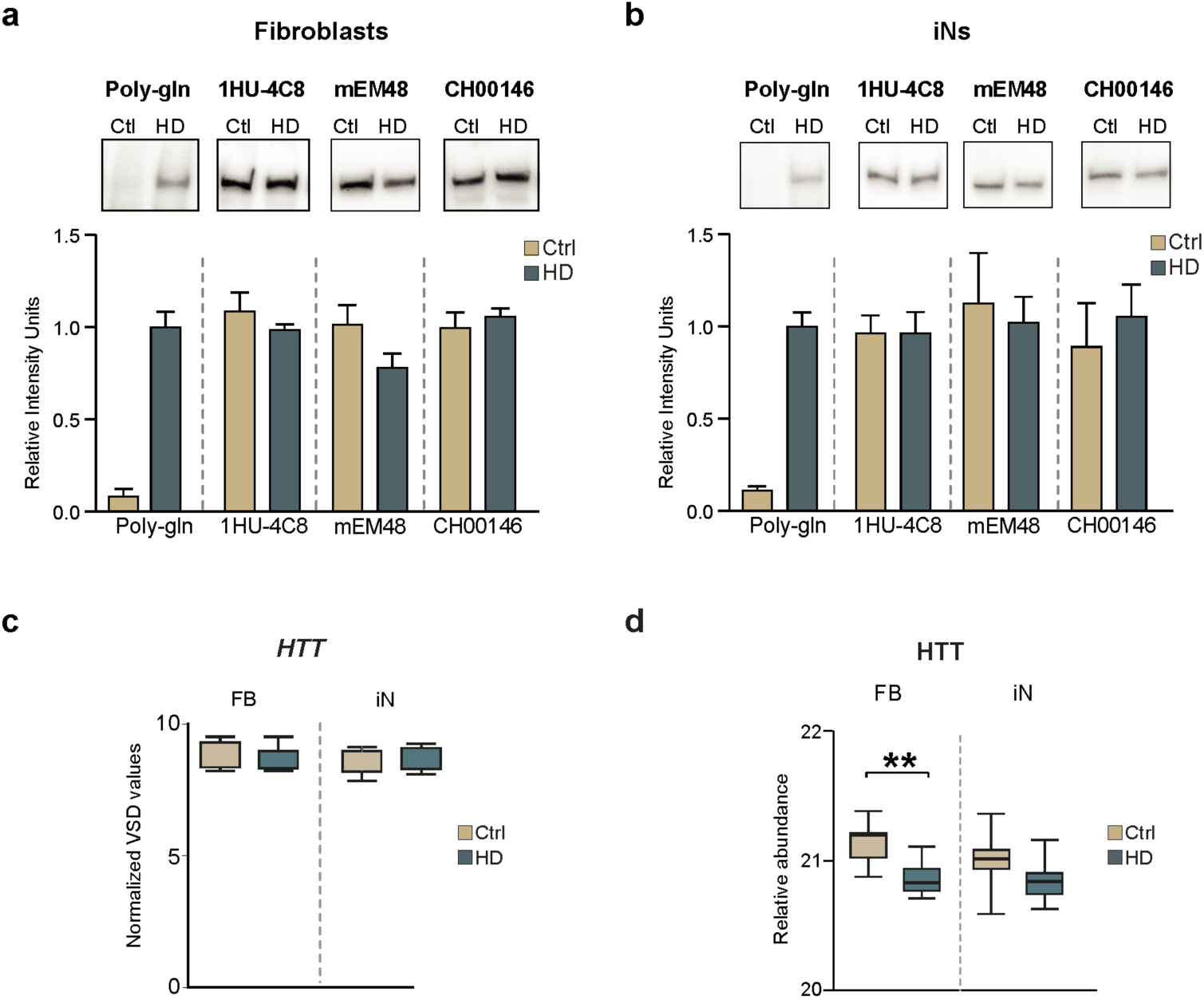
Expression of HTT protein in fibroblasts and iNs without protein aggregation. (a) Western blot quantification and representative immunoblots of HTT levels in fibroblasts (n = 3) and iNs (n = 3). (b) *HTT* RNA normalized VSD counts from control and HD fibroblasts and iNs (n = 7 control and 7 HD fibroblast and iN lines). (c) Protein abundance of HTT in control and HD fibroblasts and iNs (n = 7 control and 7 HD fibroblasts and iN lines). (p**<0.01; two-tailed unpaired T-tests were used) All data are shown as mean ± SEM. All data are shown as min/max box plots in b and c. WB values are normalized relative intensity units corrected to total protein values.

**SFigure 7.**
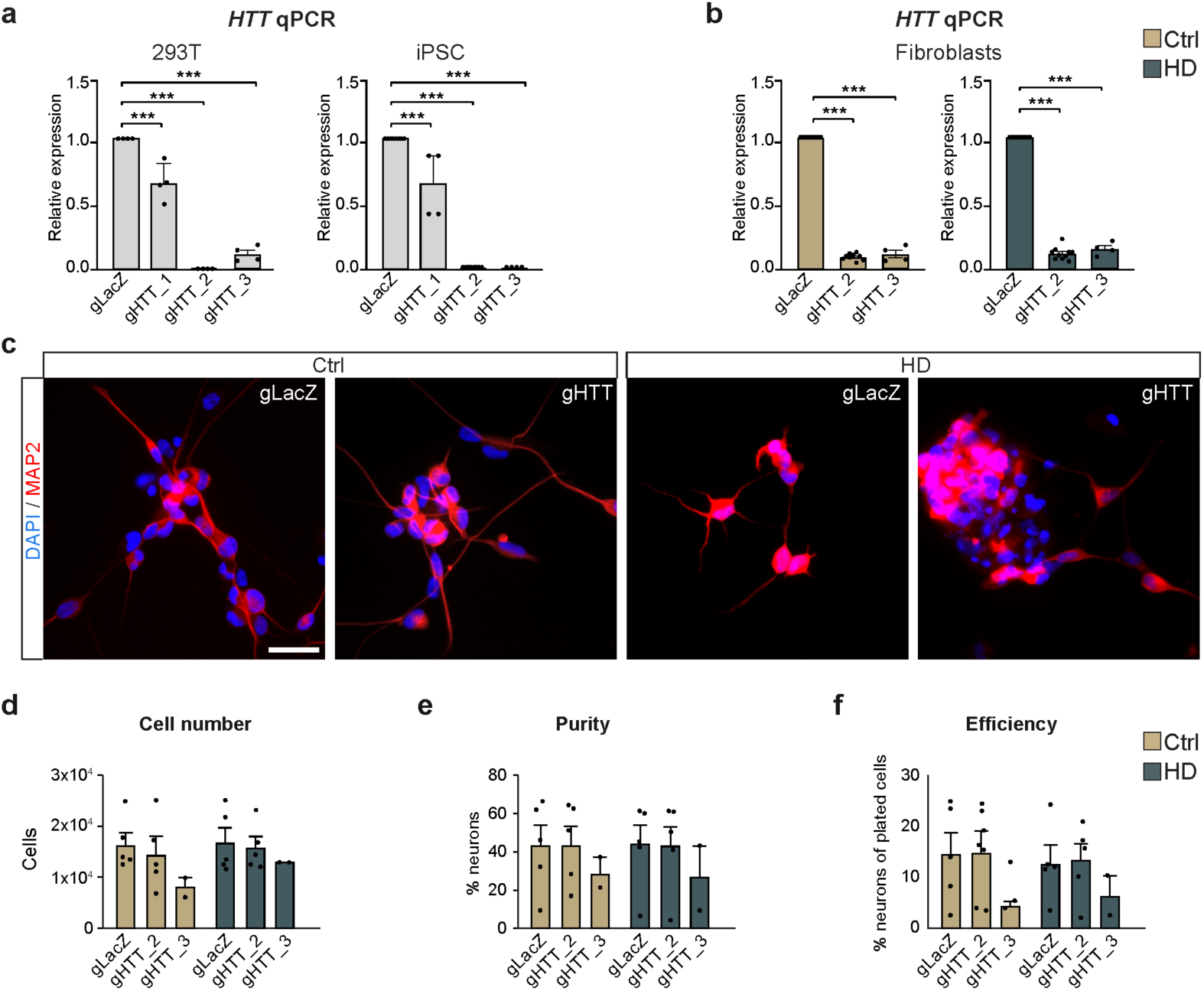
Efficient silencing of HTT using CRISPRi. Related to Figure 5. (a) 293T and iPS qRT-PCR representing silencing efficiency of the *HTT* gene using three gRNAs and vectors with MOI 10 (n = 4 for 293T cells and n = 4 for iPSc). (b) qRT-PCR representing silencing efficiency of the *HTT* gene in control and HD fibroblasts using g2RNA and g3RNA (n = 10 for LacZ and gRNA2 from 5 ctrl and 5 HD-iN lines and n = 4 for gRNA3 from 2 ctrl and 2 HD-iN lines). (c) Representative images of *HTT* and *LacZ* silencing MAP2^+^ control and HD-iN cells using CRISPRi. (d) Average DAPI^+^ cell numbers analyzed with high-content automated microscopy in *LacZ* and *HTT* g2RNA and g3RNA transduced control and HD-iNs (n = 5 ctrl and HD-iN lines for LacZ and gRNA2, n = 2 ctrl and HD-iN lines for gRNA3). (e, f) Purity and conversion efficiency of *HTT* and *LacZ* g2RNA and g3RNA transduced control and HD-iNs (n = 5 ctrl and HD-iN lines for LacZ and gRNA2, n = 2 ctrl and HD-iN lines for gRNA3). (***p<0.001; **<0.01; *p<0.05; Ordinary one-way ANOVA was used in a and b. Two-way ANOVA was used in d-f.) All data are shown as mean ± SEM. Scale bar is 50 μm.

**SFigure 8.**
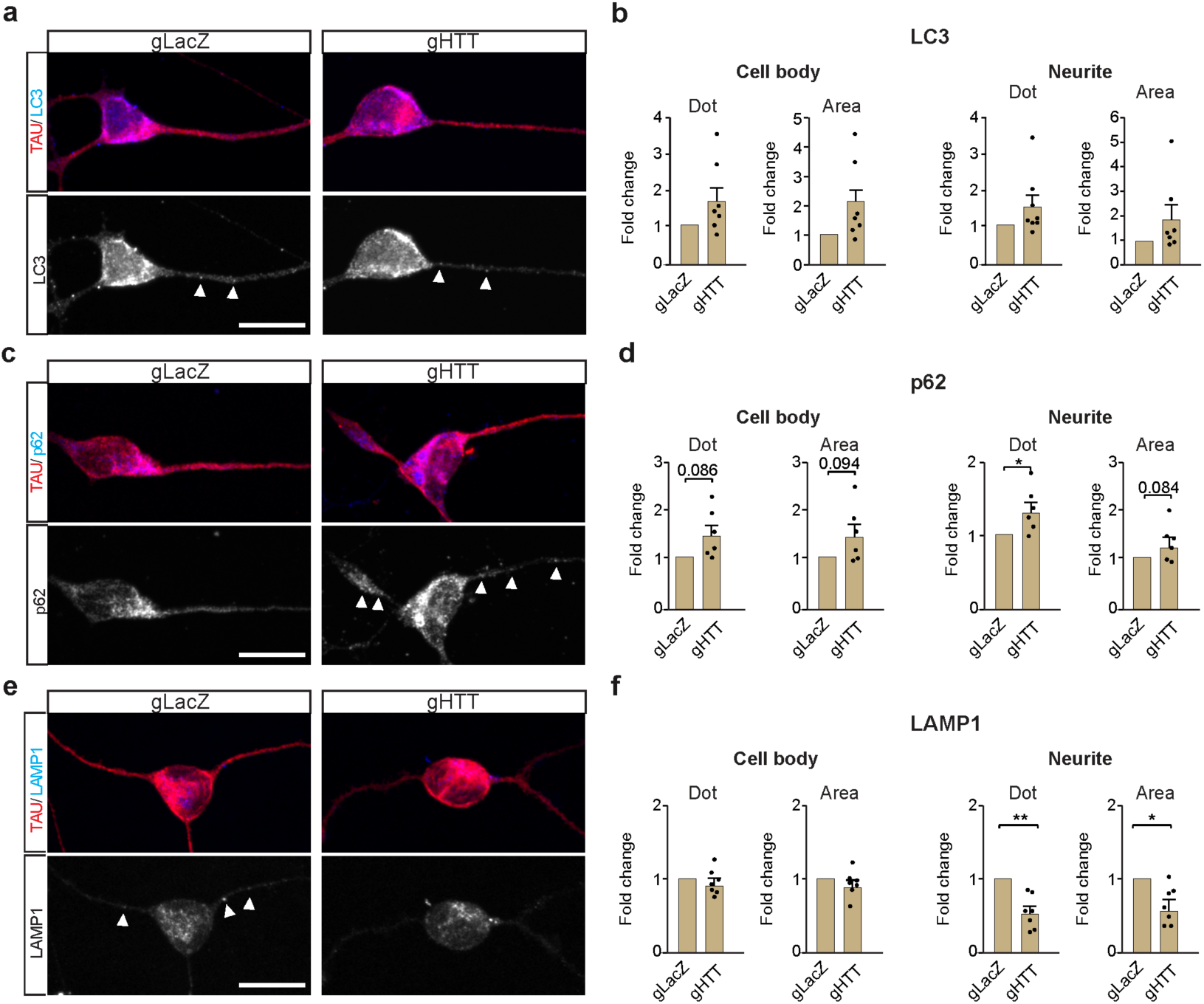
Related to Figure 5. (a-f) Representative images and statistical analysis of LC3, p62 and LAMP1 dot number and area in TAU^+^ cells in control iNs stably expressing LacZ and HTT gRNAs using CRISPRi (n = 7 replicates from 5 ctrl and 5 HD-iN lines pooling gRNA2 and gRNA3 data). (*p<0.05; Two-tailed paired t-tests were used.) All data are shown as mean ± SEM. Fold changes are presented in all graphs. Scale bar is 25 μm.

**SFigure 9.**
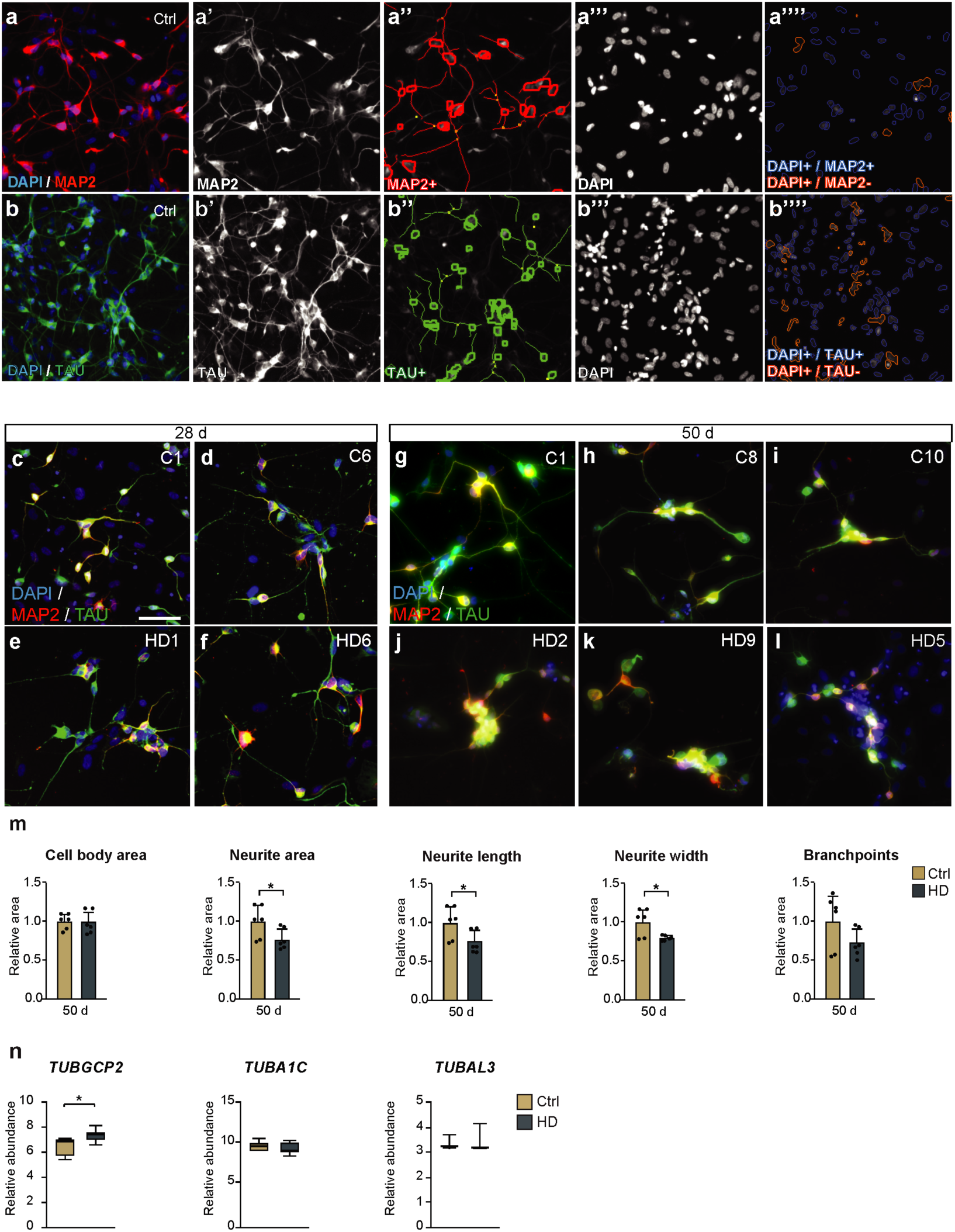
Related to Figure 6. (a) Neuronal profiling image analysis by high-content automated microscopy. DAPI^+^ cells are defined by intensity and area criteria. Border objects are excluded. MAP2^+^ or TAU^+^ cell bodies are defined by intensity measurements and area and shape on a cell-by-cell basis. Blue circles around the nuclei define the valid MAP2^+^ or TAU^+^ iNs, while orange circled nuclei are not positive for MAP2 or TAU and therefore are not counted as converted neurons. Red circles and neurites represent MAP2^+^ iNs while green circles and neurites represent TAU^+^ iNs. (c-l) Representative ICC images from control and HD patient iNs after 28 and 50 days of conversion. (m) Neuronal morphology measurement comparing control and HD-iNs after 50 days of conversion (n = 6 replicates from 3 ctrl and 3 HD-iN lines) (n) RNA expression of three different tubulin genes in control and HD-iNs (n = 7 ctrl and 7 HD-iN lines). (*p<0.05; two-tailed unpaired T-tests were used) All data are shown as mean ± SEM in m. All data are shown as min/max box plots in n. Scale bar is 50 μm.

